# Modeling methyl-sensitive transcription factor motifs with an expanded epigenetic alphabet

**DOI:** 10.1101/043794

**Authors:** Coby Viner, Charles A. Ishak, James Johnson, Nicolas J. Walker, Hui Shi, Marcela K. Sjöberg-Herrera, Shu Yi Shen, Santana M. Lardo, David J. Adams, Anne C. Ferguson-Smith, Daniel D. De Carvalho, Sarah J. Hainer, Timothy L. Bailey, Michael M. Hoffman

## Abstract

Transcription factors bind DNA in specific sequence contexts. In addition to distinguishing one nucleobase from another, some transcription factors can distinguish between unmodified and modified bases. Current models of transcription factor binding tend not take DNA modifications into account, while the recent few that do often have limitations. This makes a comprehensive and accurate profiling of transcription factor affinities difficult.

Here, we developed methods to identify transcription factor binding sites in modified DNA. Our models expand the standard A/C/G/T DNA alphabet to include cytosine modifications. We developed Cytomod to create modified genomic sequences and enhanced the Multiple EM for Motif Elicitation (MEME) Suite by adding the capacity to handle custom alphabets. We adapted the well-established position weight matrix (PWM) model of transcription factor binding affinity to this expanded DNA alphabet.

Using these methods, we identified modification-sensitive transcription factor binding motifs. We confirmed established binding preferences, such as the preference of ZFP57 and C/EBPβ for methylated motifs and the preference of c-Myc for unmethylated E-box motifs. Using known binding preferences to tune model parameters, we discovered novel modified motifs for a wide array of transcription factors. Finally, we validated predicted binding preferences of OCT4 using cleavage under targets and release using nuclease (CUT&RUN) experiments across conventional, methylation-, and hydroxymethylation-enriched sequences. Our approach readily extends to other DNA modifications. As more genome-wide single-base resolution modification data becomes available, we expect that our method will yield insights into altered transcription factor binding affinities across many different modifications.

## Introduction

Different cell types in one organism exhibit distinct gene expression profiles, despite sharing the same genomic sequence. Epigenomic regulation is essential for this phenomenon and contributes to the maintenance of cellular identity. In that regard, covalent DNA cytosine modifications have an important role in gene regulation in a number of eukaryotic species, including mice and humans.^1^ The best-studied cytosine modification is 5-methylcytosine (5mC), which entails the addition of a methyl group to the 5’ carbon of cytosine. Widely known for its effect on gene expression, 5mC occurs in diverse genomic contexts.^2,3^

Active demethylation of methylcytosine to its unmodified form proceeds through successive oxidation to 5hmC, 5fC, and 5-carboxylmethylcytosine (5caC),^4,5^ mediated by TET enzymes^6^ (Figure 1). While 5hmC has less genome-wide abundance than 5mC, it is nonetheless recognized as a stable modification.^7^ Furthermore, 5hmC is increasingly implicated in gene regulation processes.^8^ We know less about 5fC and 5caC, largely because they are even less abundant than 5hmC.^9^

**Figure 1.**
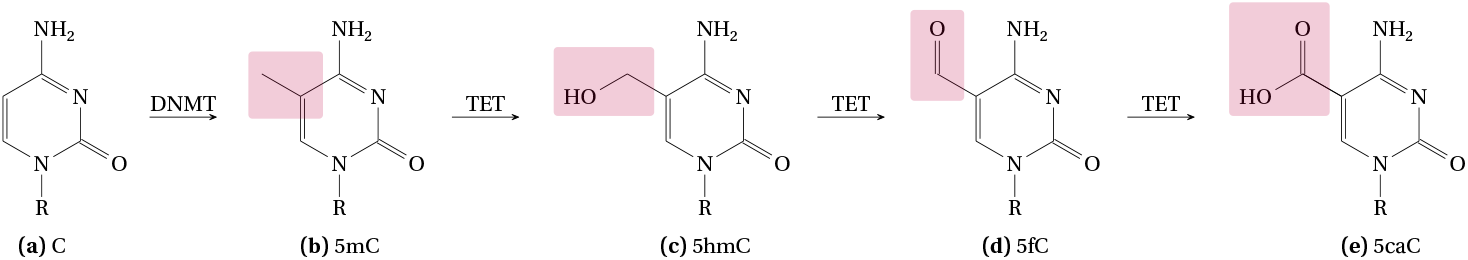
Stepwise epigenetic modification of cytosine. **(a)** DNA methyltransferase (DNMT) methylates cytosine to create **(b)** 5mC, which ten-eleven translocation (TET) enzymes oxidize to create **(c)** 5-hydroxymethylcytosine (5hmC), which TET again oxidizes to create **(d)** 5-formylmethylcytosine (5fC). Finally, TET can further oxidize 5fC to **(e)** 5-carboxylmethylcytosine (5caC), which can then return to an unmodified cytosine through decarboxylation or thymine DNA glycosylase (TDG) mediated excision, followed by base excision repair. R indicates deoxyribose and the rest of a DNA molecule. Purple rectangles indicate functional groups changed in the reaction.

In mouse embryonic stem cells (mESCs), 5fC accounts for only 0.0014 % of cytosine bases,^10^ while 5caC accounts for a miniscule 0.000335 %^4^ compared to nearly 3 % of 5mC^4^ and 0.055 % of 5hmC.^10^ Hence, fewer studies investigate the genome-wide distribution and functions of 5fC and 5caC.^11–13^ In fact, 5fC and 5caC are often regarded as mere intermediates of the demethylation cascade. Nevertheless, while it remains uncertain if 5fC and 5caC do play a distinctive and pan-tissue regulatory function, several lines of evidence suggest that they too can modulate gene expression.^8^

Although these covalent cytosine modifications do not alter DNA base pairing, they do protrude into the major and minor grooves of DNA and impact other aspects of DNA conformation.^14^ These effects can influence the DNA binding of transcription factors.^15,16^ Many transcription factors prefer specific motifs, enabling the sequence specificity of transcriptional control.^17^ The position weight matrix (PWM) model allows the computational identification of transcription factor binding sites by characterizing the position-specific preference of a transcription factor over the A/C/G/T DNA alphabet.^18^

Just as transcription factors distinguish one unmodified nucleobase from another, some transcription factors distinguish between unmodified and modified bases. For example, some transcription factors, such as MeCP2, bind to methyl-CpG.^19^ This type of non-sequence-specific modified nucleobase binding, however, occurs only in specific protein families.^20^

A few transcription factors have well-characterized modification preferences. For example, both C/EBPα and C/EBPβ have increased binding activity in the presence of central CpG methylation, formylation, or carboxylation of their canonical binding motif (consensus: `TTGC`|`GCAA`). Both DNA strands contribute and hemi-modification leads to a reduced effect.^21^ 5hmC inhibits binding of C/EBPβ, but not C/EBPα.^21^ ZFP57 also prefers methylated motifs, specifically in the context of a completely centrally-methylated `TGCCGC(R)` heptamer (red indicates methylation on the positive strand and blue on the negative strand).^22,23^

Additional methylation often occurs in ZFP57 motifs with a final guanine residue as the core binding site.^23^ Crystallography and fluorescence polarization analyses further confirm this preference.^24^ ZFP57 has successively decreasing affinity for the oxidized forms of 5mC.^24^ In contrast, the basic helix-loop-helix (bHLH) family transcription factor c-Myc, has a strong preference for unmethylated E-box motifs, often preferring the fully unmethylated CACGTG hexamer.^25,26^ Many other bHLH transcription factors also demonstrate such a preference.^27–31^

Other transcription factors also have methylation sensitivity.^32^ Protein binding microarray data demonstrated that central CpG-methylated motifs have strong binding activity for multiple transcription factors.^15^ Interestingly, these data also show that these motifs often differ from the unmethylated sequences that those transcription factors usually bind. Some transcription factors may even show increased binding in the presence of 5caC.^33^ In *Arabidopsis thaliana,* among 327 transcription factors, 248 (76 %) exhibited sensitivity to covalent DNA modifications, with 14 preferring modified DNA.^34^

Transcription factors act as both readers and effectors of methylation.^20^ They may bind to a modified base to prevent its modification, or, in some instances, to increase the likelihood of its modification. Alternatively, transcription factors could bind to reverse an existing modification. These scenarios could occur in different genic contexts, potentially mediated by different motif groups. Even factors within the same family may have differences in modified binding preferences, conferring additional specificity or assisting in stable protein-DNA complex formation. This regulatory interplay^20,35^ highlights the need for additional genome-wide characterizations of transcription factor binding preferences in the context of modified DNA.

The role of modified DNA in transcription factor binding has motivated the development of a computational framework to elucidate and characterize altered motifs. A comprehensive *in vitro* analysis, coupled with selected follow-up crystal structures, revealed the mechanistic basis for some 5mC interactions.^36^ A random forest^37,38^ combined genomic and methylation data^39^ to predict transcription factor binding. Those predictions, however, did not attempt to predict the preference of factors for methylated DNA.^39^ The MethMotif database enumerates methylated transcription factor motifs.^40^

Most recently, Grau et al.^41^ employed an expanded alphabet genome from whole genome bisulfite sequencing (WGBS) data, for 5mC only. Their focus differs from ours, however. They emphasize that their models go beyond PWMs, the standard model to describe transcription factor DNA binding specificities and allow for intra-motif dependencies. Their comparisons mainly focus on classification performance bench-marking in differentiating bound versus unbound sequences. Song et al.^42^ demonstrate an *in vitro* method to assess modification-specific preferences of all cytosine states. They demonstrate distinct preferences of both symmetric and hemi-modifications. Most recently, Hernandez-Corchado et al.^43^ developed a joint model of accessibility and methylation sequence. They used this model to explore a large number of chromatin immunoprecipitation-sequencing (ChIP-seq) datasets, assessing many transcription factor binding site preferences for 5mC.

Existing work has often indirectly analyzed the impact of modified bases on binding, focused on improved motif elucidation itself, or often categorized modified binding preferences in a largely binary fashion. Mostly, when modeling the affinity of transcription factors for DNA sequences, previous work has not treated modified nucleobases as first-class objects akin to unmodified nucleobases, adding artificial distinctions unlikely to reflect the underlying biophysical interactions. There has been a dearth of large-scale comprehensive analyses including modified motifs. Also, there has been an absence of specific experimental follow-up to predicted motif preferences, directly detecting modified bases.

Here, we describe methods to analyze covalent DNA modifications and their effects on transcription factor binding sites by introducing an expanded epigenetic DNA alphabet. While others proposed expanding the genomic alphabet in other ways,^44^ we (in our earlier preprint of this work)^45^ and Ngo et al.^46,47^ first proposed expanding it in this context for facilitating bioinformatic analyses of cytosine modifications. Unlike our work, however, Ngo et al.^46^ focused on motif identification in this expanded alphabet. We, rather, leverage existing tools to focus on downstream consequences, such as distinct groups of modified-preferring motifs and specific predictions of modified binding preferences. We introduce Cytomod, a software to integrate DNA modification information into a single genomic sequence and we detail the use of extensions to the Multiple EM for Motif Elicitation (MEME) Suite^48^ to analyze 5mC and 5hmC transcription factor binding site sensitivities. We validate our predictions for the transcription factor OCT4 by providing conjoint cleavage under targets and release using nuclease (CUT&RUN)^49,50^ datasets across conventional, methylated-, and hydroxymethylated-enriched sequences. Our results especially highlight that most factors can bind in both unmodified and modified contexts, to varying extents and often with different groups of motifs. While it was previously known that DNA methylation affects binding, here we show that modified motifs are considerably more complex than previously appreciated, and that many new motifs with varied modified binding preferences exist, to different extents across a variety of transcription factors.

## Results

### Expanded-alphabet genomes facilitate the analysis of modified base data

We created an expanded-alphabet genome sequence using oxidative (ox) and conventional WGBS maps of 5mC and 5hmC for naive *ex vivo* mouse CD4^+^ T cells.^51^ We expand the standard A/C/G/T alphabet, adding the symbols `m` (5mC), `h` (5hmC), `f` (5fC), and `c` (5caC). We also designed and implemented symbols for the reverse strand, preserving information of complements (Methods). This allows us to more easily adapt existing computational methods, that work on a discrete alphabet, to work with epigenetic cytosine modification data.

Next, we generated individual modified genomes across four replicates of combined ox and conventional WGBS data^51^ and for a variety of modified base calling thresholds. These calibrated modified genomes allowed us to accurately assess transcription factor binding site affinities, for both 5mC and 5hmC. In order to construct modified genome sequences, specific to the varied epigenetic state of a cell type, we designed the Cytomod software. It allows us to rapidly construct combined threshold-specific modified genomes, using single-base resolution data. Modified genomes with our expanded alphabet allowed us to deploy our methods across large datasets including those from Encyclopedia of DNA Elements (ENCODE).^52^

We used these modified genome sequences as the basis for the extraction of genomic regions implicated by ChIP-seq data for all assessed transcription factors. Using the thresholds discovered in the murine analyses, we created conventional 5mC maps for the human K562 erythroid leukemia cell line,^53,54^ using ENCODE WGBS data.

We updated the MEME Suite^48^ and associated software to work with custom alphabets, such as our expanded epigenomic alphabet. We created the MEME::Alphabet Perl module as part of the implementation. Others can use this module to rapidly obtain suitable expanded-alphabet definitions, making it easier to extend older code bases. These changes allow comprehensive analyses of epigenetic states, including their impacts on transcription factor binding, with support for any additional modified bases. Furthermore the software improvements make all future MEME Suite tools compatible with expanded alphabets, enabling continuing innovation and insights in these areas.

### Our methods yield suitable base-calling thresholds for downstream analyses

The grid search for transcription factor binding thresholds at 0.01 increments allowed us to determine suitable thresholds (0.3 and 0.7) for further investigation (Figure S1). Overall, this grid search demonstrated the suitability of a wide range of thresholds, indicating the range for which one can adequately maintain both specificity and sensitivity of modified binding detection. For example, *de novo* analyses of C/EBPβ confirmed the preference for methylated DNA, with methylated motifs having much greater central enrichment than their unmethylated counterparts, at both the 0.3 (Figure 3) and 0.7 thresholds (Figure 4). The bounding of suitable thresholds provided by the grid search analysis will likely prove useful for assessing future datasets as well.

**Figure 2.**
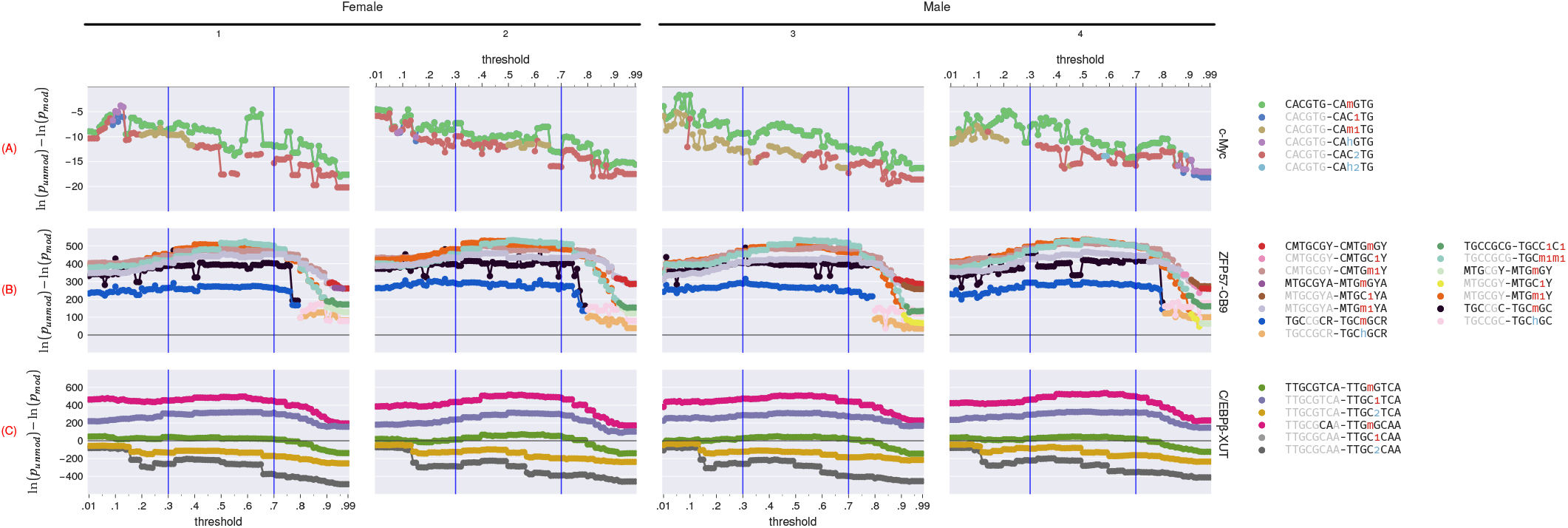
Relationship between unmodified versus modified motif statistical significance of central enrichment (from CentriMo^55^) and modified base calling thresholds across different WGBS and oxWGBS specimens, in mice.^51^. We compare each unmodified motif, at each threshold, to its top three most significant modifications for c-Myc and C/EBPβ, but only the single most significant modification for ZFP57. The displayed motif pairs changes at individual thresholds, depending on which motif pairs stay in the top three. See Table 1 and Table 3 for an overview of modified base notation. Sign of value indicates preference for the unmodified (negative) motif or the modified (positive) motif. Rows: single ChIP-seq replicates for a particular transcription factor target, one each of: (**A**) c-Myc (mESCs; Krepelova et al.^58^), (**B**) ZFP57 (CB9 mESCs; Strogantsev et al.^23^), and (**C**) C/EBPβ (C2C12 cells;ENCFF001XUT). Columns: replicates of WGBS and oxWGBS (mouse CD4^+^ T cells;Kazachenka et al.^51^).

**Figure 3.**
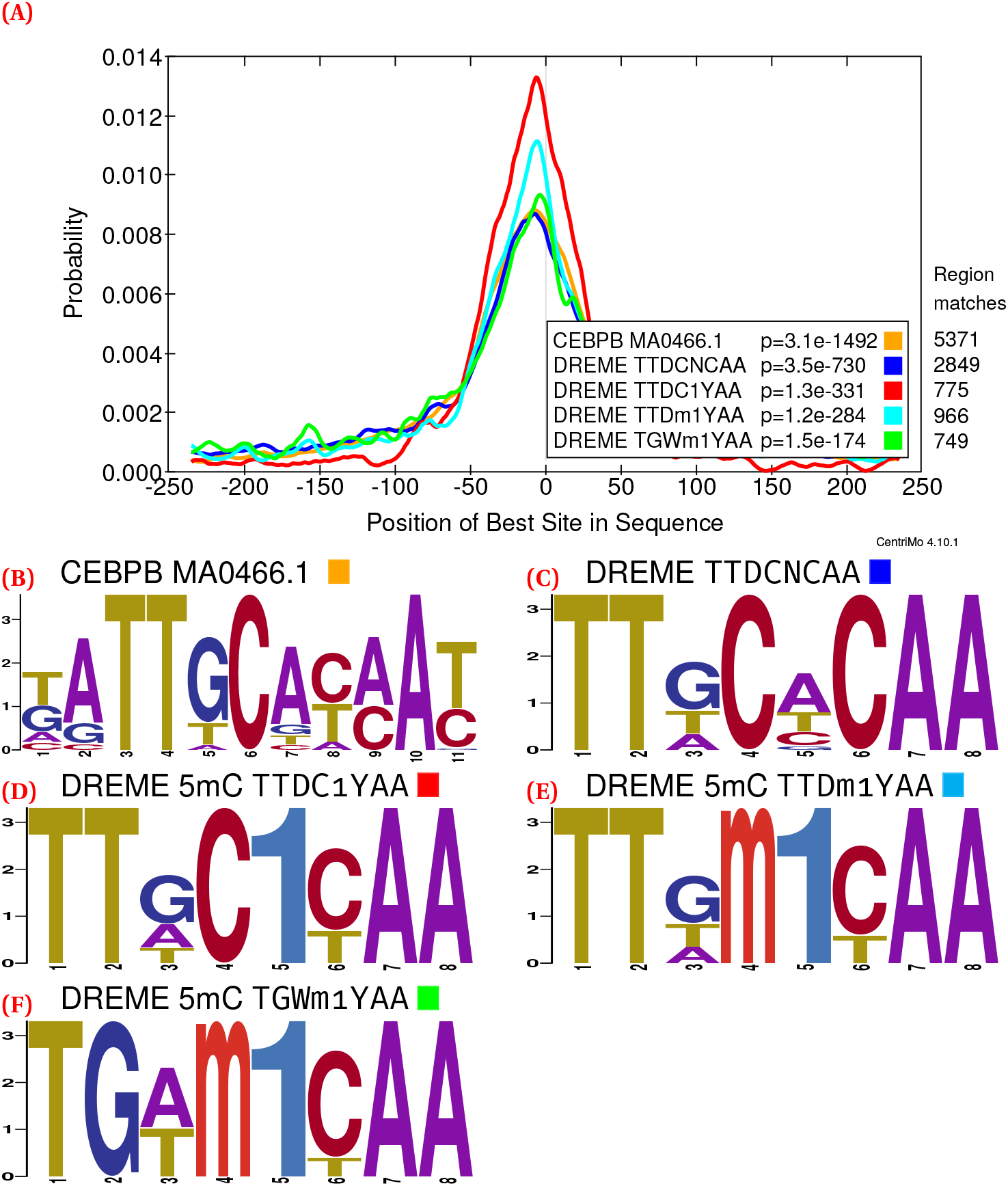
C/EBPβ (GSM915179; 11 434 ChIP-seq peaks) CentriMo analysis of *de novo* and JASPAR motifs (Methods). Depicts female replicate 2 of the combined WGBS and oxWGBS data^51^ at a 0.3 modification threshold. (**A**) the CentriMo result with the JASPAR C/EBP β motif (orange), top Discriminative Regular Expression Motif Elicitation (DREME) unmethylated C/EBPβ motif (blue), and DREME methylated motifs (red, cyan, and green). (**B**) Sequence logo of the JASPAR C/EBPβ motif. (**C**) Sequence logo of the top DREME unmodified motif. (**D**) Sequence logo of the top DREME methylated motif. (**E**) Sequence logo of the second DREME methylated motif. (**F**) Sequence logo of the third DREME methylated motif. Listed p-values computed by CentriMo.^55^ For consistency, we depict the JASPAR sequence logo using MEME’s relative entropy calculation and colouring.

**Figure 4.**
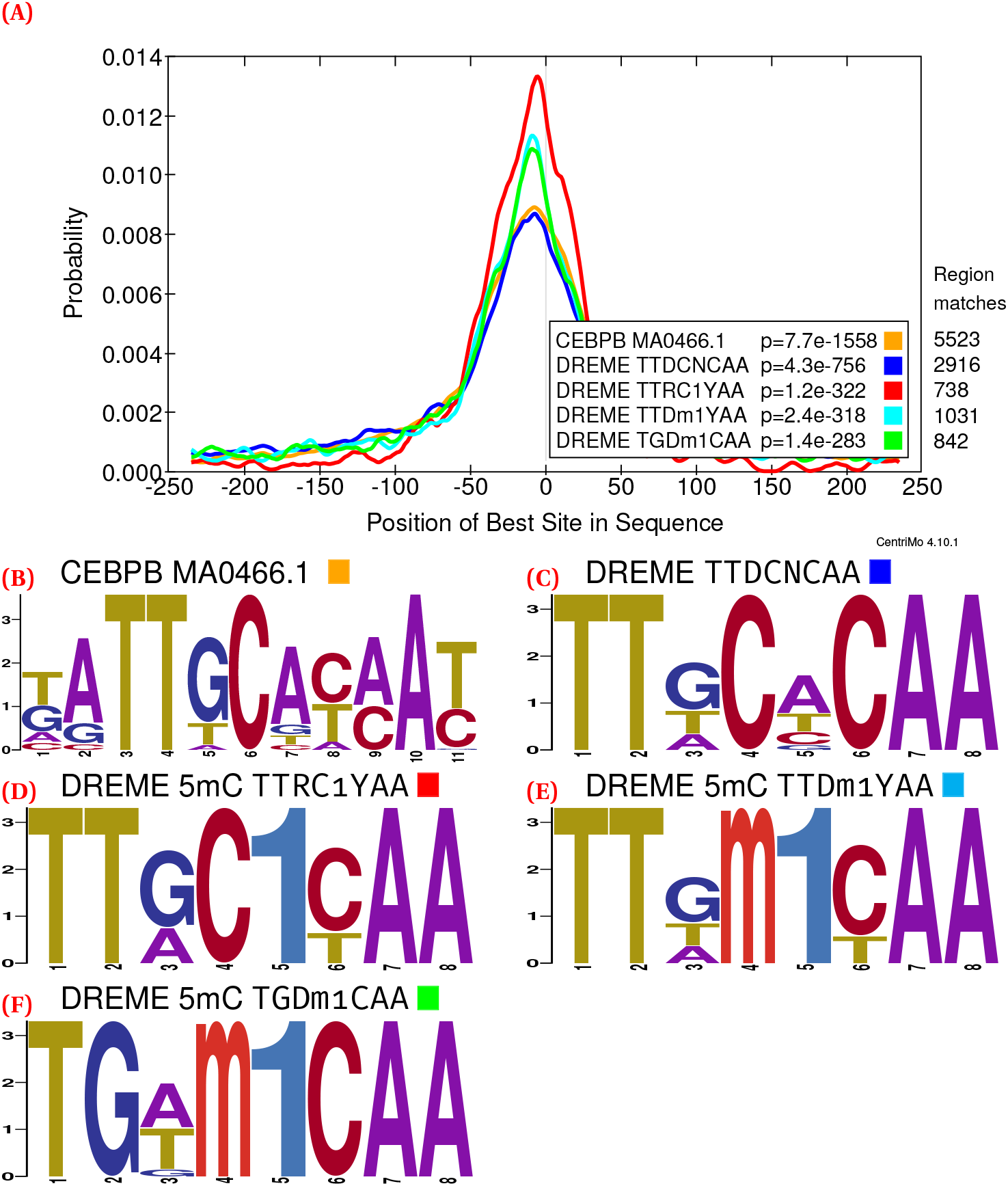
C/EBPβ (GSM915179; 11 434 ChIP-seq peaks) CentriMo analysis of *de novo* and JASPAR motifs (Methods). Depicts female replicate 2 of the combined WGBS and oxWGBS data^51^ at a 0.7 modification threshold. (**A**) the CentriMo result with the JASPAR C/EBP β motif (orange), top DREME unmethylated C/EBPβ motif (blue), and DREME methylated motifs (red, cyan, and green). (**B**) Sequence logo of the JASPAR C/EBPβ motif. (**C**) Sequence logo of the top DREME unmodified motif. (**D**) Sequence logo of the top DREME methylated motif. (**E**) Sequence logo of the second DREME methylated motif. (**F**) Sequence logo of the third DREME methylated motif. Listed p-values computed by CentriMo.^55^ For consistency, we depict the JASPAR sequence logo using MEME’s relative entropy calculation and colouring.

### Hypothesis testing reveals altered transcription factor binding preferences

#### Expanded-alphabet analysis shows results consistent with known preferences

We used a hypothesis testing approach on the expanded-alphabet sequence to examine the preferences of transcription factors for modified or unmodified DNA. First, we analyzed three transcription factors with previously known methylation or hydroxymethylation sensitivities. ZFP57^23^ and C/EBPβ^21^ show a preference for methylated DNA, while c-Myc prefers unmethylated DNA.^25,26^ Additionally, C/EBPβ has reduced affinity for hydroxymethylated DNA.^21^

We used the known preferences as controls to calibrate our modification-calling thresholds, and to validate our approach. We used c-Myc as the positive control for an unmethylated binding preference.^25,26^ As positive controls for methylated binding preferences, we used both ZFP57 and C/EBPβ^21–24^ (Methods).

In this hypothesis testing framework, we tested all known unmodified transcription factor binding motifs against all possible 5mC and 5hmC modifications at all CpG dinucleotides. That is, for each unmodified and modified version of all motifs, across every transcription factor, we assessed the motif’s expected DNA binding affinity using the adjusted central enrichment p-value from CentriMo^55^ (Methods). For this analysis, we included motifs of interest from *de novo* results, and we partially or fully changed the base at a given motif position to each modified base, to comprehensively assess its affinity (Table 3; Methods). To compare all binding affinities, we subtracted the log_10_-transformed p-value of the modified motif from the unmodified motif. Positive values for this difference represented a preference for the modified motif, while negative values represented a preference for the unmodified.

**Table 1.** The expanded epigenetic alphabet. This includes the known modifications to cytosine and symbols for each guanine complementary to a modified nucleobase.

| Covalent Cytosine Modification |  |  | Complement |  |
| --- | --- | --- | --- | --- |
| Abbreviation | Name | Symbol | Name | Symbol |
| 5mC | 5-Methylcytosine | **m** | Guanine:5-Methylcytosine | **1** |
| 5hmC | 5-Hydroxymethylcytosine | **h** | Guanine:5-Hydroxymethylcytosine | **2** |
| 5fC | 5-Formylcytosine | **f** | Guanine:5-Formylcytosine | **3** |
| 5caC | 5-Carboxylcytosine | **c** | Guanine:5-Carboxylcytosine | **4** |

**Table 2.**
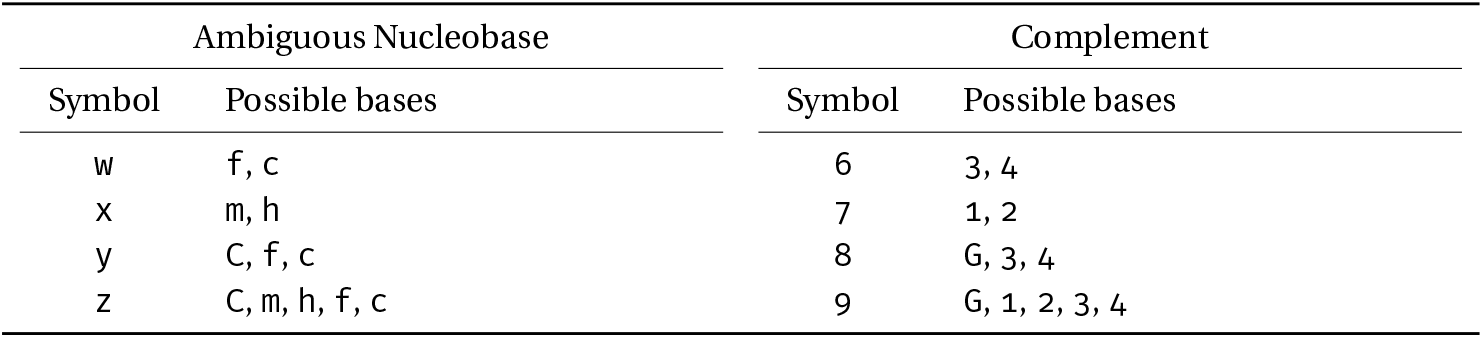
Ambiguous bases for uncertain modification states. The MEME Suite recognizes these ambiguity codes in the same manner as the ambiguous bases already in common usage, such as R for A or G in the conventional DNA alphabet.

| Ambiguous Nucleobase |  | Complement |  |
| --- | --- | --- | --- |
| Symbol | Possible bases | Symbol | Possible bases |
| w | f, c | 6 | 3, 4 |
| x | m, h | 7 | 1, 2 |
| y | C, f, c | 8 | G, 3, 4 |
| z | C, m, h, f, c | 9 | G, 1, 2, 3, 4 |

**Table 3.** Illustrative examples of possible changes made to convert unmodified motifs to specific modified counterparts, for downstream hypothesis testing. We use stacked letters like simple sequence logos. At these positions, `N` represents any base frequencies other than the base being modified. These make up the other positions in the motif’s PWM. 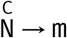 indicates that a position containing cytosine is modified by setting all base frequencies other than `m` to 0 and setting the frequency of `m` to 1. Conversely, 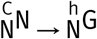 indicates that a position containing cytosine is modified by replacing the frequency apportioned to `C` with `h`, leaving the other base frequencies at that position unmodified. We portray the second base of each dinucleotide as having a frequency of 1. This second base, however, could also comprise different bases of various frequencies, including the base shown.

| Modification description | Unmodified motif | Modified motif |
| --- | --- | --- |
| Full CpG modification to 5mC | $\begin{smallmatrix} \text{C} \\ \text{N} \end{smallmatrix} \text{G}$ | m1 |
| Full CpG modification to 5hmC | $\begin{smallmatrix} \text{C} \\ \text{N} \end{smallmatrix} \text{G}$ | h2 |
| Partial CpG modification to 5hmC | $\begin{smallmatrix} \text{C} \\ \text{N} \end{smallmatrix} \text{G}$ | $\begin{smallmatrix} \text{h} \\ \text{N} \end{smallmatrix} 2$ |
| Partial CpG hemi-modification to 5hmC | $\begin{smallmatrix} \text{C} \\ \text{N} \end{smallmatrix} \text{G}$ | $\begin{smallmatrix} \text{h} \\ \text{N} \end{smallmatrix} \text{G}$ |
| Full CpT modification to either 5mC or 5hmC | $\begin{smallmatrix} \text{C} \\ \text{N} \end{smallmatrix} \text{T}$ | xT |

The expected transcription factor binding preferences for c-Myc, ZFP57, and C/EBPβ held across all four biological replicates of WGBS and oxWGBS data and for all investigated modified nucleobase calling thresholds (Figure 2). The thresholds we investigated, representing modification confidence, varied from 0.01–0.99 inclusive, at 0.01 increments. We also obtained the same results for multiple different ChIP-seq replicates for these three transcription factors (Figure S1). Perturbations of binding assessments, such as peak-calling stringency (Figure S2) and required degree of motif statistical significance (Figure S3) demonstrated the robustness of our results.

None of the c-Myc log p-value differences exceeded zero, confirming that c-Myc favours unmodified E-box motifs over modified c-Myc motifs. Two methylated motifs had the greatest increase in predicted binding affinity for C/EBPβ: TTGmGCAA and TTGC1TCA (see Table 1 and Table 3 for an overview of modified base notation). As expected, ZFP57 favours binding to modified nucleobases over their unmodified counterparts. The well-known `TGCmimi` motif^23^ had one of the greatest increases in predicted binding affinity of ZFP57 for modified DNA.

While ZFP57 had a strong preference for methylated DNA, we also observed a noticeable preference for hydroxymethylated DNA (Figure 5D). CentriMo quantifies these preferences,^55^ both in terms of p-value significance, and in terms of the centrality of motif concentration (Methods).

**Figure 5.**
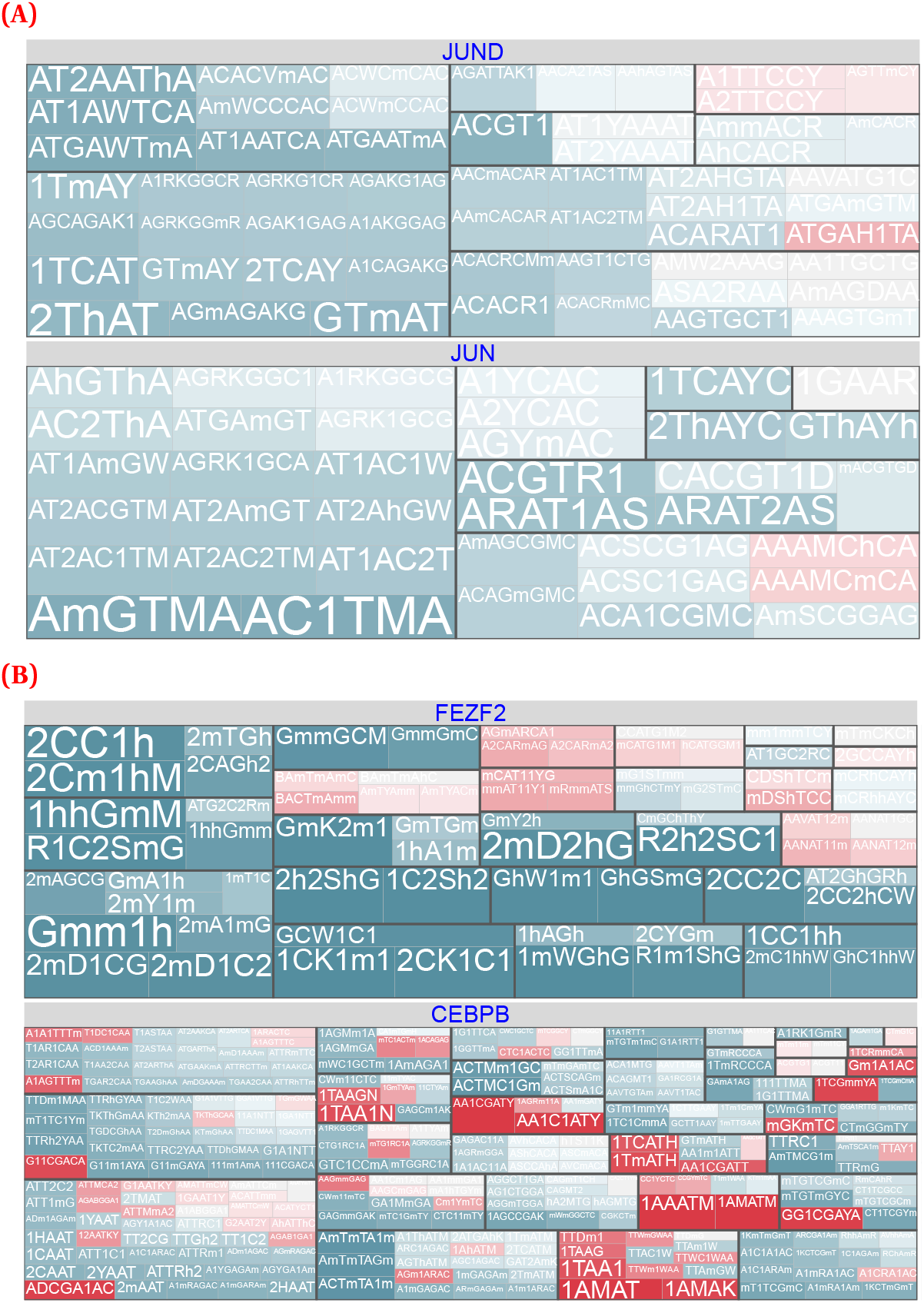
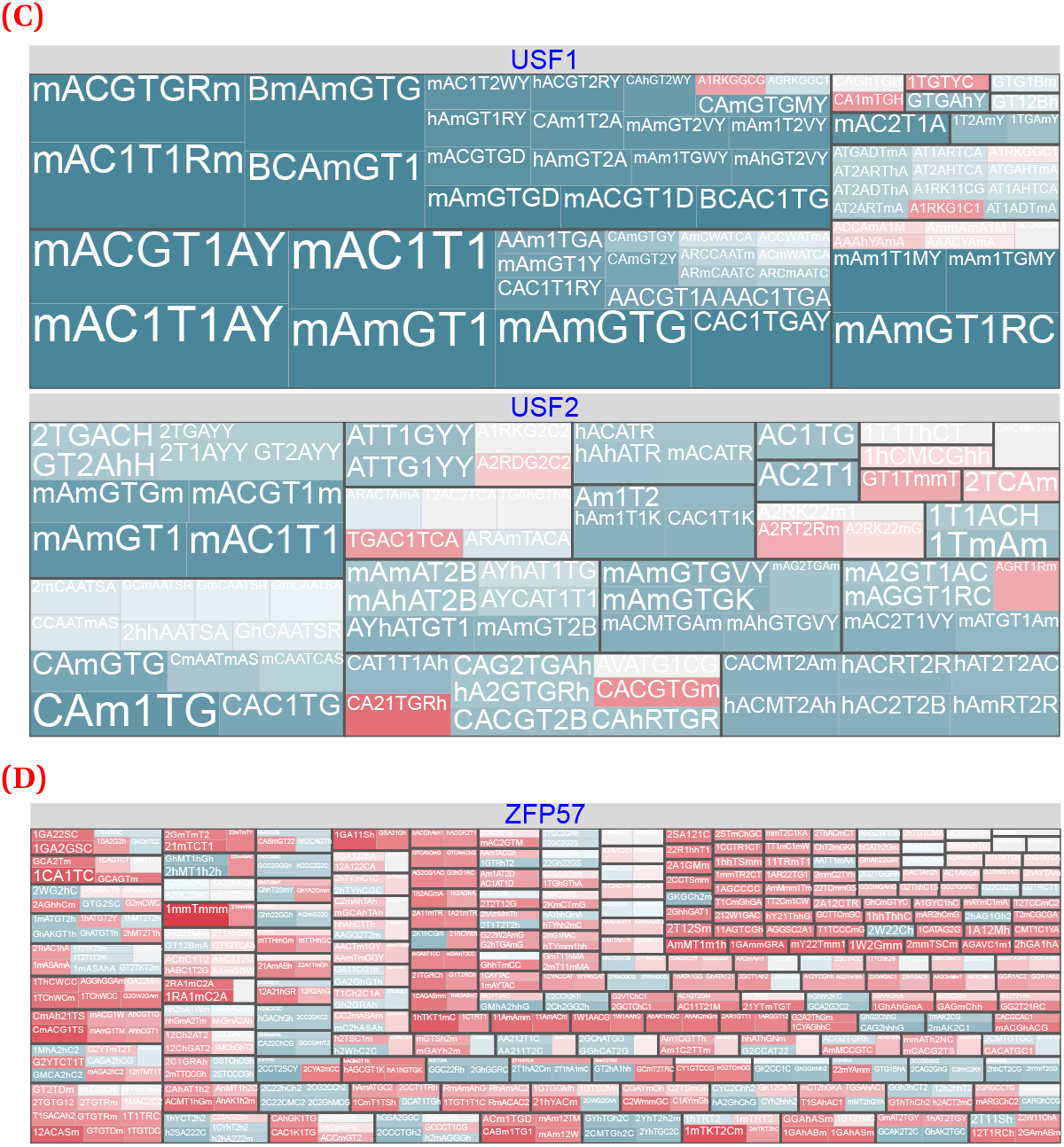
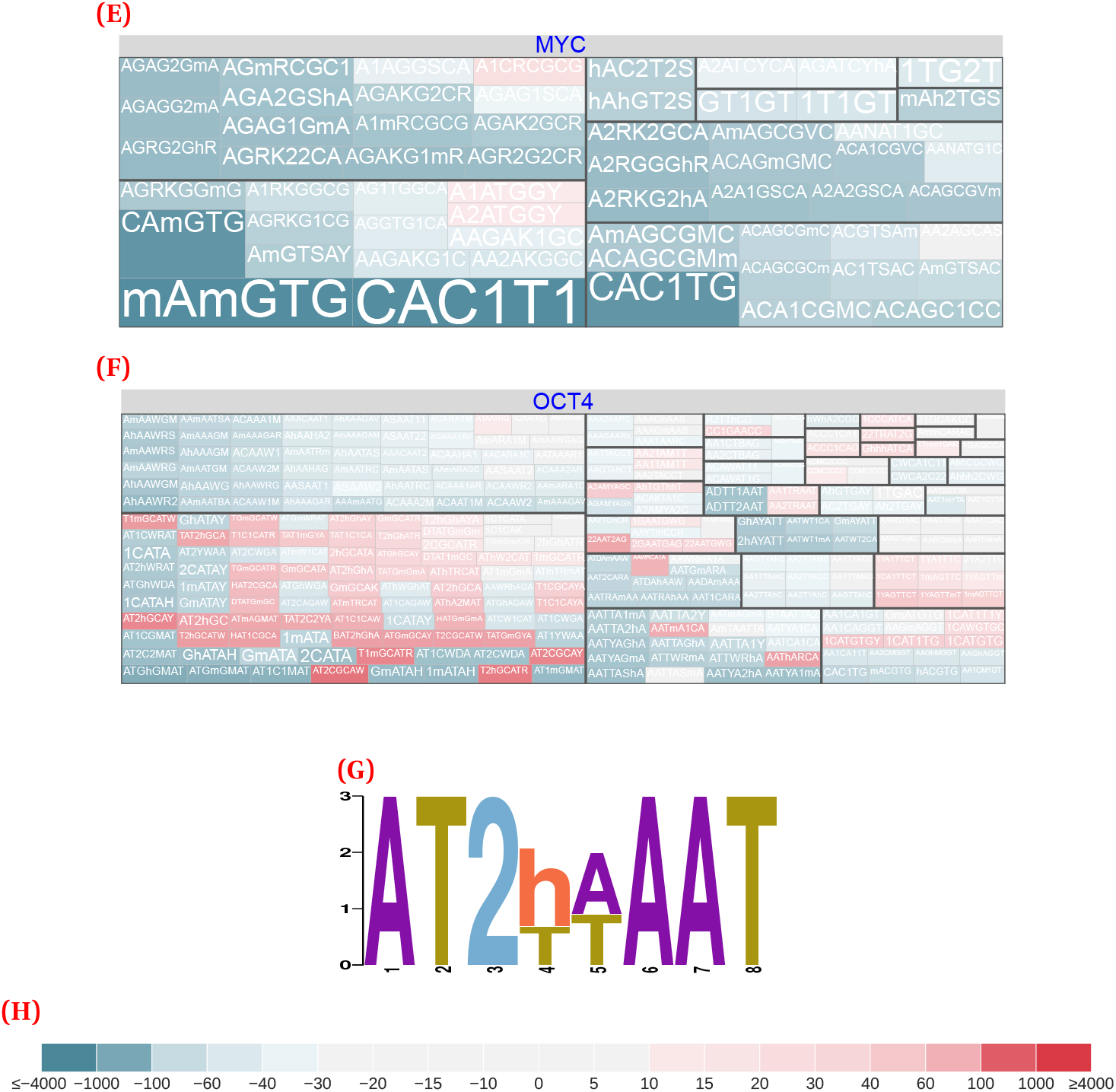
Modified versus unmodified motifs, combining score and cluster information, for selected transcription factors. These plots come from non-spike-in calibrated data, for the 500 bp regions surrounding peak summits. We clip scores above or below ±4000 and plot them at threshold to maximize dynamic range where most scores occur. Some combinations of the displayed hypothesis pairs had multiple data points (for example, multiple identical hypothesis pairs, but for different data sub-types or stringencies). We aggregated these data points bylplotting the maximum score. An asymmetric, diverging, colour scale further highlights modification-preferring motifs. We depict a larger selection of transcription factors in Figure S6.(**A**) JUND and JUN. (**B**) PEZF2 and C/EBPβ. (**C**) USF1 and USF2. (**D**) ZFP57. (**E**) c-Myc. (**F**) OCT4. (**G**) The most highly!significant and centrally enriched DREME motif, for OCT4 hmC-Seal CUT&RUN in mESCs (replicate 1). See Table 1 and Table 3 for an overview of modified base notation. (**H**) Colour scale.

CentriMo reported 328 of 393 total ZFP57 methylated motifs with a score >0 (median: 171.2; max: 2292, exemplifying the strong preference) across all our assessed ZFP57 datasets. This included motifs from mESCs, from both our previously-mentioned BC8/CB9 Strogantsev et al.^23^ datasets, and 2 motifs, both scoring positively, from Quenneville et al.^22^ (Figure 5D). Hydroxymethylated CpGs had a substantially smaller increase in binding affinity than methylated motifs (Figure 2), but still greater than the completely unmethylated motif. CentriMo reported 291 of 435 total ZFP57 hydroxymethylated motifs with a score >0 (median: 152.4; max: 1379) across all motifs in our BC8/CB9 datasets (Figure 5D). Our Quenneville et al.^22^ analysis did not reveal any sufficiently significant hydroxymethylated motifs of any score. Most modified motifs that scored above zero, however, had at least one 5mC and one 5hmC nucleobase (Figure 5D, red motifs). Therefore, our expanded-alphabet methodology recapitulates the observation that ZFP57 has the greatest binding affinity for motifs containing 5mC, followed by 5hmC, and then by unmodified cytosine.^24^

Overall, we these positive control results for known binding preferences allowed us to tune model parameters, permitting confident broader application of our methods. This led us to discover novel modified motifs, across the wider array of transcription factors to follow.

#### Expanded-alphabet analysis enables comparisons across a wide array of transcription factors

Similarities in protein structure of transcription factors might form a useful categorical framework for expectations regarding modified base affinity. To that end, we looked for shared preferences among families of transcription factors for modified or unmodified bases in both the mouse and the human data. Mostly, we defined families with TFClass^56,57^ (Methods).

We and others^27–31^ have found a consistent preference for unmethylated binding motifs across a broad selection of bHLH transcription factors. Some motifs for bHLH transcription factors had a putatively modified preference versus their unmodified JASPAR counterparts. Unmodified *de novo* motifs we generated for the same transcription factors, however, consistently had more significant p-values (Figure S6). This suggests that, as expected, these transcription factors usually preferred the unmodified motif. The leucine-zipper subfamily of bHLH transcription factors, however, had a subset of motifs that preferred to bind in a modified context. For example, both USF1 and USF2 preferred to bind in unmodified and modified contexts, to differing extents, and had mixed binding preferences within motif clusters (Figure 5C; Discussion).

Many zinc finger family motifs displayed a propensity toward modified motifs, but not all. EGR1/ZIF268/NGFI-A, a Cis_2_–His_2_ zinc finger, showed a moderate binding preference for methylated DNA, with multiple positively-scoring hypothesis pairs, including some >100. These very high scores indicate an exceptionally strong predicted binding preference for the modified, over the unmodified, nucleobases.

Conversely, ZNF384/CIZ/TNRC1 of the same sub-family, present in both our mouse and human analyses, had only weak evidence of a preference for binding modified DNA, with only a single hypothesis pair scoring above 10 (Figure S1). We suspect this factor intrinsically has the ability to bind in both unmodified and modified contexts, perhaps with a weak modified preference. This would likely hold across quite different tissue types. Unlike most of our analyzed transcription factors, this result occurred at a significant level in both our mouse and human datasets, allowing us to form this more general conclusion.

While we can use our methods to analyze and group transcription factors by their families, few clear signals of strong preferences nicely stratify in this manner. This suggests that complex preferences tend to outweigh family-specific patterns. Factors such as local epigenetic state or tissue type likely play a larger role in locus-specific transcription factor binding.

### Our hypothesis testing confirms C/EBPβ’s dichotomous binding preferences

C/EBPβ provides an excellent test case for the impact of modified bases on transcription factor binding because of its dichotomous preferences for 5mC versus 5hmC.^21^ Our method recapitulated this preference, across all ChIP-seq datasets, for all replicates of ox and conventional WGBS. Methylated motif pairs generally had positive ratios, whereas hydroxymethylated motif pairs had negative ratios (Figure S3).

One positive strand, hemi-methylated motif (`TTGmGTCA`) presented an exceptional case. Surprisingly, we observed a preference for the unmodified motif over its hemi-methylated motif. Unlike the consensus C/EBPβ motif, this motif corresponds to the chimeric C/EBP|CRE octamer. This chimeric transcription factor has a more modest preference toward its methylated DNA motif.^21^ Nonetheless, we would still have expected a weak preference for the hemi-methylated motif, over its unmodified counterpart. Additionally, we found greater enrichment for hemi-methylation than complete methylation, which contradicts findings of both strands contributing to the preferential binding of C/EBPβ.^21^ This may arise from technical issues with hemi-methylation in our modified sequence, or because our methods have greatest accuracy only within specific cell types or contexts.

### Many transcription factors bind in modified and unmodified contexts, with variable motif preferences

We analyzed 144 transcription factors to characterize their overall motifs and their affinities to methylated and hydroxymethylated DNA. Leveraging our hypothesis testing approach and normalized CentriMo-based scoring methods, we assessed all detected motifs, and specifically characterize their likely binding affinities. Our analyses revealed that several factors bind in both modified and unmodified contexts (Figure 5). Unlike prior analyses which often aimed to binarize binding preferences, our results highlight that most factors can bind in both contexts, albeit to varying extents and with different motifs.

For example, protein binding microarray analyses have led to the conclusion that binding of the transcription factor JUND is “uniformly inhibited by 5mC”.^59^ Overall, these data accord well with our results, for which almost all tested hypothesis pairs (37/44) showed an unmethylated preference.

Nevertheless, closer inspection of these prior analyses reveals that they contain a small group of motifs where JUND showed a slight preference for 5mC.^59^ Specifically, at least 8 cytosine-containing motifs have 5mC z-scores above zero, with at least 3 such motifs having scores of close to 30. Similar findings applied to 5hmC. This indicates that JUND likely has at least some preference for hydroxymethylated motifs, despite mostly preferring to bind in unmodified contexts.

In our analysis, JUND showed a preferences for binding to 7 motifs (including one hydroxymethylated motif) if the motifs were methylated (Figure 5A). JUN, a transcription factor related to JUND, showed similar preferences (Figure 5A).

FEZF2 appeared to have two completely different motifs, with no overlap in their preferences for modified versus unmodified cytosine (Figure 5B). Indeed, removing modification information from modified-preferring FEZF2 motifs led to a single motif cluster, distinct from the main unmodified motif clusters. Therefore, two distinct motif classes for FEZF2 appear to exist.

Many of the motifs we found, across a wide array of transcriptions factors including those discussed above, were novel. Many motif groups, especially when viewed as collapsed or root motifs, often have similarity with previously reported motifs. Nonetheless, within these motif groups, we often find additional variations, as well as a number of entirely new motif groups, for most transcription factors.

More than half of the transcription factors we assessed bound almost or entirely exclusively in unmodified contexts (Figure S6). Specifically, if we limit our analysis to transcription factors without even a single slightly positive motif, 49.3% of factors had all tested hypothesis pairs score below zero. This varied across our overall dataset, with other factors having some occasional modified preferences.

### Modified-base CUT&RUN validates our predictions for OCT4

While OCT4 bound to a number of motifs in an unmodified context, some OCT4 motifs preferred binding in both methylated and hydroxymethylated states. A preference of OCT4 for methylated motifs has previously been reported,^36^ but we are unaware of any reports of a preference of OCT4 for hydroxymethylated sequences. Interestingly, those hydroxymethylated motifs appeared to predominantly cluster either on their own, or with the canonical OCT4 homo- and hetero-dimer motifs, rather than mixing with other motif groups, such as those belonging to methylated motifs or co-factors.

We validated our OCT4 predictions, by performing CUT&RUN^49,50^ experiments in mESCs, with conventional, bisulfite-converted, and hmC-Seal-seq library preparations. These three sets of library preparations allowed us to characterize the modification states of OCT4-bound fragments across both methylated and hydroxymethylated contexts.

We observed that OCT4 has a strong preference to bind in a hydroxymethylated context, in line with our predictions. When comparing unconverted, conventional CUT&RUN to hmC-Seal-seq CUT&RUN, OCT4 and similar *de novo* motifs were preferentially bound in hydroxymethylated context (Figure 6; top DREME motif: *p* = 4.5 × 10^-149^; top OCT4 motif: *p* = 2.5 × 10^-13^) than in the unmodified context (top DREME motif: *p* = 4.1 × 10^-104^; top OCT4 motif: *p* = 1.1 × 10^-8^), all with comparable or greater motif centrality. These motifs also had at least similar preferences for binding in a methylated context (Figure S5).

**Figure 6.**
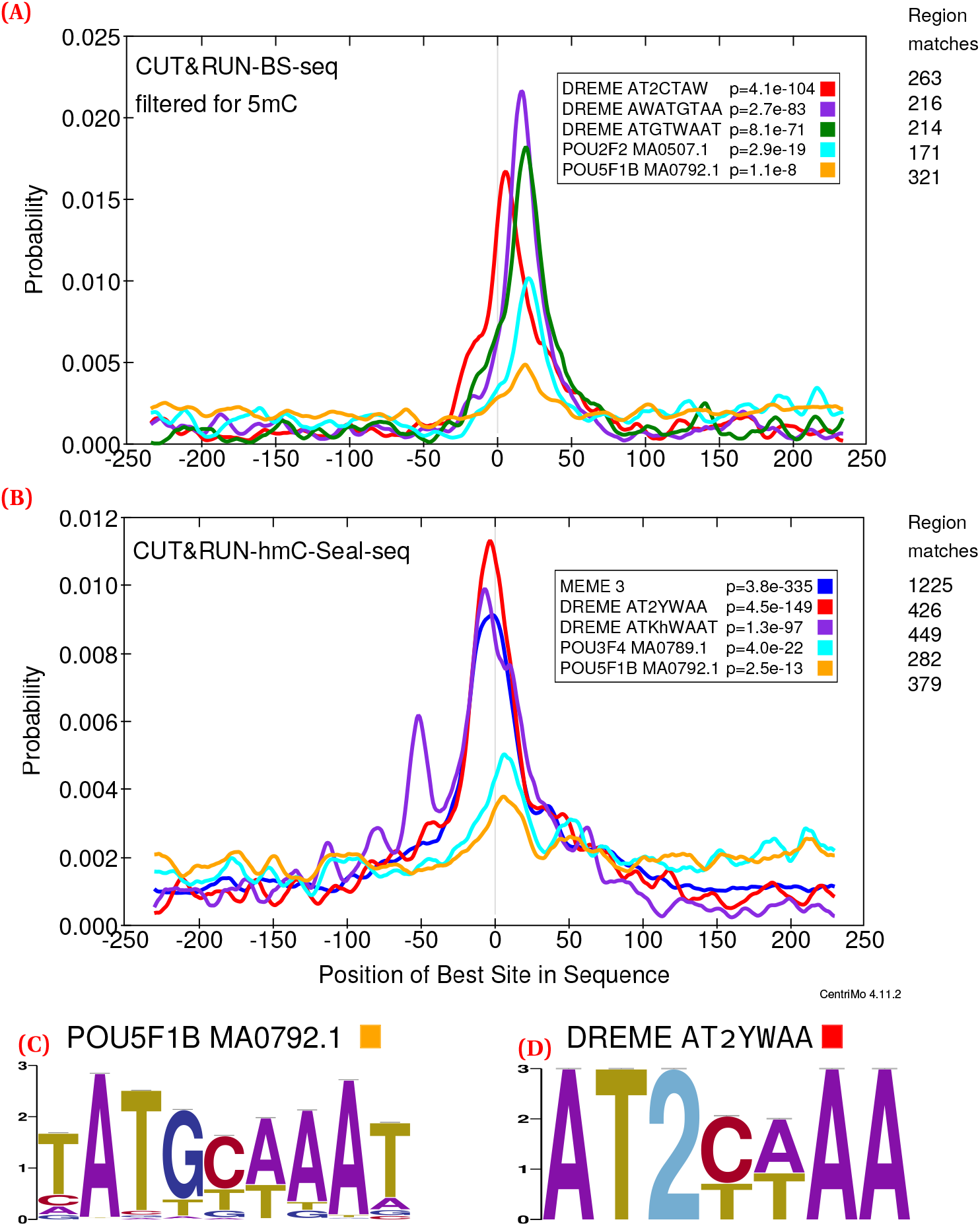
CentriMo^55^ results for replicate 1 of OCT4 CUT&RUN in mESCs. Motifs include the top three DREME motifs with colour indicating rank: first (red);second (purple), third, where applicable (dark green). Motifs also include the top non-POU5F1 JASPAR motif (cyan), and the JASPAR POU5F1B motif (orange). Both of these motifs come from the JASPAR 2020^60^ core vertebrate set. We generated these results using 500 bp regions centred upon the summits of MACS 2^61^ peaks generated from those CUT&RUN fragments ≤120 bp. We called peaks using IgG controls and without any spike-in calibration (Methods). Listed p-values computed by CentriMo.^55^ For consistency, we depict the JASPAR sequence logo using MEME’s relative entropy calculation and colouring. (**A**) bisulfite-converted (methylated;2797 CUT&RUN peaks) sequences. (**B**) hmC-Seal (3974 CUT&RUN peaks) sequences. Also depicts the top MEME^62^ *de novo* motif (blue). (**C**) Sequence logo of the POU5F1B motif (MA0792.1). (**D**) Sequence logo of the top DREME *de novo* motif.

The predicted cluster that included the canonical POU5F1B JASPAR motif (MA0792.1) also showed enrichment for 5hmC motifs. Overall, our findings suggest that OCT4 specifically binds hydroxymethylated nucleobases, in concert with methylated and unmodified binding sites.

## Methods

### An expanded epigenetic alphabet

To analyze DNA modifications’ effects upon transcription factor binding, we developed a model of genome sequence that expands the standard A/C/G/T alphabet. Our model adds the symbols `m` (5mC), `h` (5hmC), `f` (5fC), and `c` (5caC). This allows us to more easily adapt existing computational methods, that work on a discrete alphabet, to work with epigenetic cytosine modification data.

Each symbol represents a base pair in addition to a single nucleotide, implicitly encoding a complementarity relation. Accordingly, we add four symbols to represent G when paired with modified C: `1` (G:5mC), `2` (G:5hmC), `3` (G:5fC), and `4` (G:5caC) (Table 1). This ensures that complementation remains a lossless operation. The presence of a modification alters the base pairing properties of a complementary guanine,^14^ which this also captures. We number these symbols in the same order in which the TET enzyme acts on 5mC and its oxidized derivatives (Figure 1).^5^

Many cytosine modification-detection assays only yield incomplete information of a cytosine’s modification state. For example, conventional bisulfite sequencing alone determines modification of cytosine bases to either 5mC or 5hmC, but cannot resolve between those two modifications.^5^ Even with sufficient sequencing to disambiguate all modifications, we require statistical methods to infer each modification from the data, resulting in additional uncertainty. To capture common instances of modification state uncertainty, we also introduce ambiguity codes: `z/9` for a cytosine of (completely) unknown modification state, `y/8` for a neither hydroxymethylated nor methylated cytosine, `x/7` for a hydroxymethylated or methylated cytosine, and `w/6` for a formylated or carboxylated cytosine (Table 2). These codes are analogous to those defined by the Nomenclature Committee of the International Union of Biochemistry already in common usage, such as for unknown purines (R) or pyrimidines (Y).^63,64^

### Cytomod: method for creation of an expanded-alphabet genome sequence

Like most epigenomic data, abundance and distribution of cytosine modifications varies by cell-type. Therefore, we require modified genomes for a particular cell-type and would not necessarily expect downstream analyses to generalize. Accordingly, we first need to construct a modified genome that pertains to the organism, assembly, and tissue type we wish to analyze. This modified genome uses the described expanded alphabet to encode cytosine modification state, using calls from single-base resolution modification data.

To do this, we created a Python program called Cytomod. It loads an unmodified assembly and then alters it using provided modification data. It relies upon Genomedata^65^ and NumPy^66^ to load and iterate over genome sequence data. Cytomod can take the intersection or union of replicates pertaining to a single modification type. It also allows one to provide a single replicate of each type, and potentially to run it multiple times to produce multiple independent replicates of modified genomes. It permits flagging of ambiguous input data, such as when only possessing conventional bisulfite sequencing data, therefore yielding only `x/7` as modified bases. Cytomod additionally produces browser extensible data (BED^)67,68^ tracks for each cytosine modification, for viewing in the University of California, Santa Cruz (UCSC)^67^ (Figure 7) or Ensembl genome browsers.^69^

**Figure 7.**
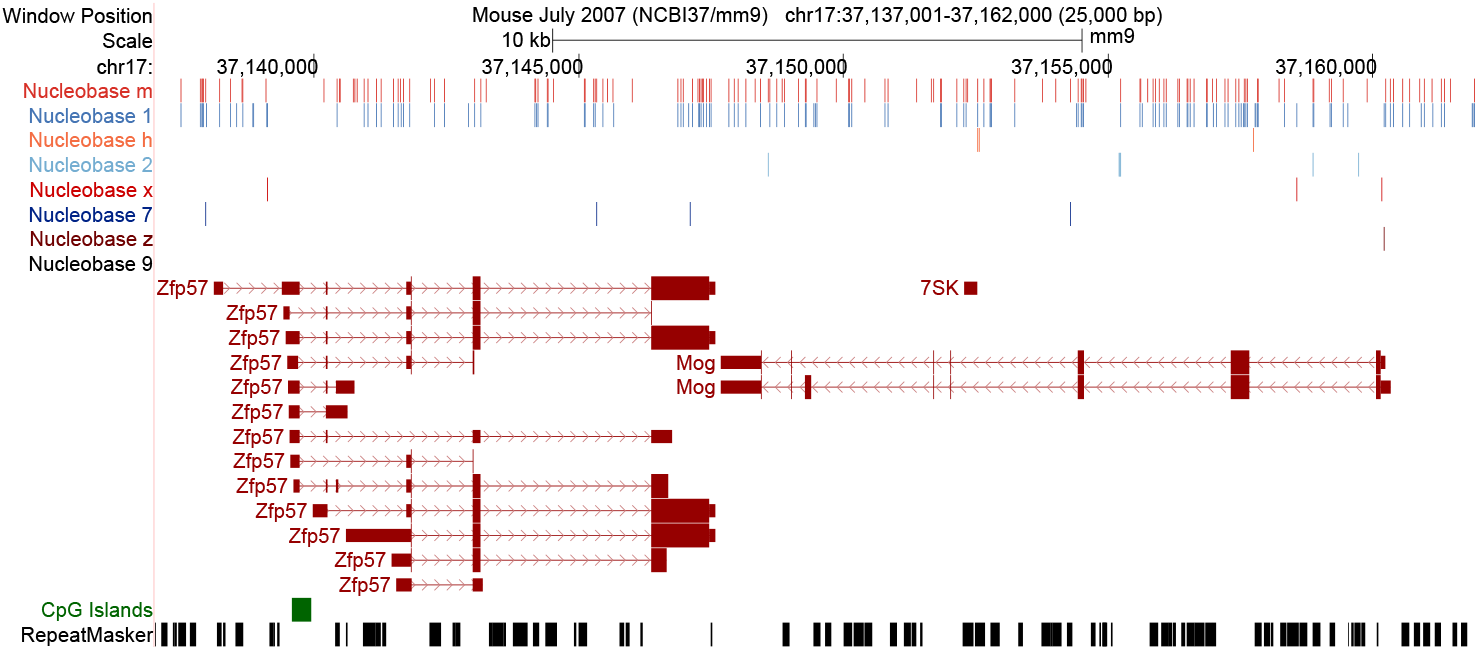
Differential cytosine modification status in naive mouse T cells for a 25 kbp region (within cytoband 17qB1) surrounding *Zfp57* and *Mog.* This UCSC Genome Browser^67^ display includes RepeatMasker^70^ regions, CpG islands,^71^ GENCODE^72^ genes, and calls for modified nucleobases `h` (5hmC), `m` (5mC), `x` (5mC/5hmC), `z` (C with unknown modification state), `1` (G:5mC), `2` (G:5hmC), `7` (G:5mC/5hmC), and `9` (G:C with unknown modification state).

We subjected the unaligned, paired-end, binary alignment/map (BAM) files output from the sequencer to a standardized internal quality check pipeline. Widely known to work well, we selected Bismark^73^ for alignment.^74,75^ We use the following processing pipeline: sort the unaligned raw BAM files in name order using Sambamba^76^ (version 0.5.4);convert the files to FASTQ,^77^ splitting each paired-end (using BEDTools^78^ version 2.23.0 `bamtofastq`);align the FASTA files to NCBI m37/mm9 using Bismark^73^ (version 0.14.3), which used Bowtie 2,^79–81^ in the default directional mode for a stranded library; sort the output aligned files by position (using Sambamba `sort`); index sorted, aligned, BAMs (SAMtools^82^ version 1.2 `index`);convert the processed BAM files into the format required by Meth-Pipe, using `to-mr`; merge sequencing lanes (using direct concatenation of `to-mr` output files) for each specimen (biological replicate), for each sex, and each of WGBS and oxidativeWGBS (oxWGBS); sort the output as described in MethPipe’s documentation (by position and then by strand);remove duplicates using MethPipe’s `duplicate-remover`; run MethPipe’s `methcounts` program;and finally run maximum likelihood methylation levels (MLML).^83^ After alignment, we excluded all random chromosomes. We use modifications called beyond a specified threshold (as described below) as input for Cytomod (with Genomedata^65^ version 1.36.dev-r0).

In bulk data, usually one considers a base modified or not using some threshold, above which one “calls” a particular modified base or set of possible modifications. There exist several ways to perform modified base calling, generally first involving computing a proportion of modification, at a specific position. We use the MLML^83^ method to do this. Then, we must decide the value sufficient to call a modification downstream.

MLML^83^ outputs maximum-likelihood estimates of the levels of 5mC, 5hmC, and C, between 0 and 1. It outputs an indicator of the number of conflicts—an estimate of methylation or hydroxymethylation levels falling outside of the confidence interval computed from the input coverage and level. An abundance of conflicts can indicate the presence of non-random error.^83^ We assign z/9 to all loci with any conflicts, regarding those loci as having unknown modification state. Our analysis pipeline accounts for cytosine modifications occurring in any genomic context. It additionally maintains the data’s strandedness, allowing analyses of hemi-modification.

### Mouse expanded-alphabet genome sequences

We used conventional and oxidative WGBS data generated for naive CD4^+^ T cells, extracted from the spleens of C57BL/6J mice, aged 6 weeks–8 weeks. The dataset authors obtained a fraction enriched in CD4^+^ T cells, by depletion of non-CD4^+^ T cells by magnetic labelling, followed by fluorescence-activated cell sorting to get the CD4^+^, CD62L^+^, CD44^low^, and CD25^-^ naive pool of T cells. We previously published these data^51^ as part of the BLUEPRINT project^84^ (GSE94674; GSE94675). We analyzed biological replicates separately, 2 of each sex.

For our mouse datasets, we aligned sequencing reads with Bowtie 2^79–81^ version 2.2.4. We used MethPipe^85^ ([development version](https://github.com/smithlabcode/methpipe), commit 3655360) to process the data.

We used our mouse datasets to calibrate our modified base calling thresholds. MLML^83^ combines the conventional and oxidative bisulfite sequencing data to yield consistent estimations of cytosine modification state. In our case, with two inputs per mouse run (WGBS and oxWGBS), we obtain values of 0, 1, or 2. We created modified genomes using a grid search, in increments of 0.01, for a threshold t, for the levels of 5mC (m) and 5hmC (h), as described in Figure 8.

**Figure 8.**
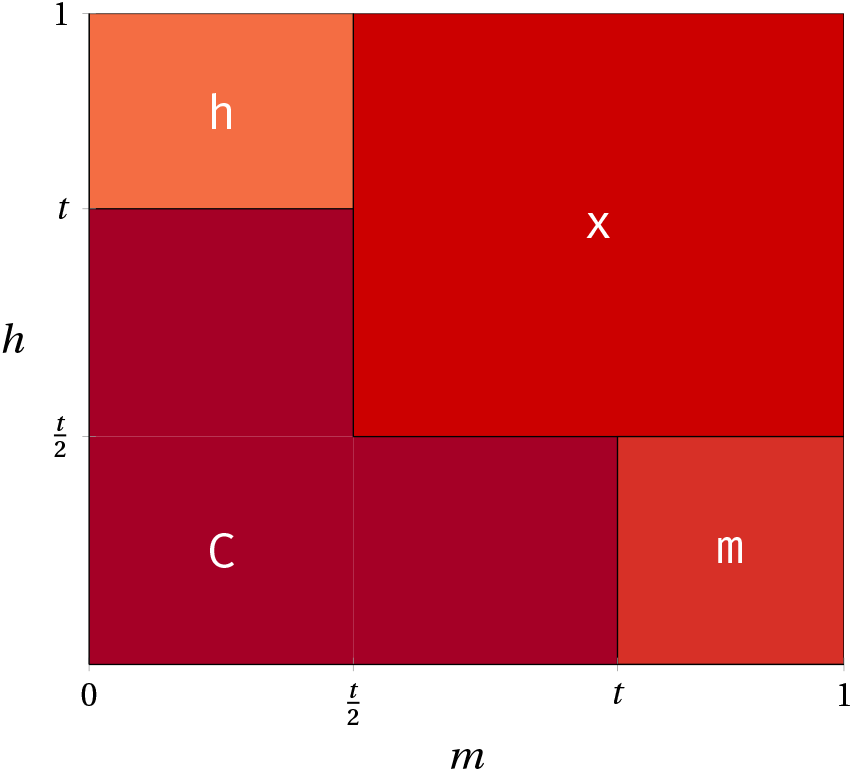
Conditions on the MLML^83^ confidence levels of 5mC (m) and 5hmC (h) in relation to a threshold t, that lead to the calling of different modified nucleobases. We call a modification if *m* or *h* equal or exceed the threshold. These base assignments assume that MLML had no conflicts for the locus under consideration. If any conflicts occur, we use `z` as the base, irrespective of the values of *m* or h. We depict bases for the positive strand only, and complement those occurring on the negative strand, as outlined in Table 1 and Table 2.

We use half of the threshold value for assignment to `x/7`, since we consider that consistent with the use of the full threshold value to call a specific modification. Namely, if *t* suffices to call 5mC or 5hmC alone, *m* + *h* ≥ t ought suffice to call `x/7`.

### Human expanded-alphabet genome sequence

Using publicly available ENCODE WGBS data (ENCFF557TER and ENCFF963XLT), we created K562 (RRID: CVCL_0004) modified genome for the GRCh38/hg38 assembly at 0.3 and 0.7 stringencies. WGBS data alone does not differentiate 5mC and 5hmC. As the data cannot differentiate between these states, one might represent them as x. Nonetheless, we represent modified bases from the WGBS data as m, both for convenience, and because, in most cases, these positions are just methylated. We processed these datasets, as previously described. We aligned human datasets with Bowtie 2^79–81^ version 2.2.3 and processed with MethPipe^85^ release version 3.4.2.

### Detection of altered transcription factor binding in modified genomic contexts

Following creation of expanded-alphabet genome sequences, we performed transcription factor binding site motif discovery, enrichment and modified-unmodified comparisons. Here, we use mouse assembly NCBI m37/mm9 for all murine analyses, since we wanted to make use of all Mouse ENCOD^E86^ ChIP-seq data (RRID: CVCL_0188) without re-alignment nor lift-over. Specifically, we used the *Mus musculus* Illumina (San Diego, CA, USA) iGenome^87^ packaging of the UCSC NCBI m37/mm9 genome. This assembly excludes all alternative haplotypes as well as all unreliably ordered, but chromosome-associated, sequences (the so-called “random” chromosomes). While ideal for downstream analyses, that assembly does not suffice for aligning data ourselves. Exclusion of these additional pseudochromosomes might deleteriously impact alignments, by resulting in the inclusion of spuriously unique reads. Therefore, we used the full UCSC NCBI m37/mm9 build when aligning to a reference sequence. For our human datasets, we used GRCh38/hg38, with all K562 ENCODE datasets.

We used all K562 peak calls, processed as outlined below, from Human ENCODE ChIP-seq data, from preliminary data processed for Karimzadeh et al.^88^. We briefly recapitulate the processing steps here. First, they align the raw reads with Bowtie 2^79–81^ (version 2.2.3). Then, they then de-duplicate reads using SAMtools^82^ (version 0.1.19) and filter for those with a mapping quality of greater than 10 using option -bq 10. Finally, they call peaks without any control, using MACS 2^61^ (version 2.1.0) `callpeak` and options `--qvalue 0.001 --format ‘BAM’ --gsize ‘hs’ --nomodel --call-summits`.

We updated the MEME Suite^48^ to work with custom alphabets, such as our expanded epigenomic alphabet. Additionally, we created the MEME::Alphabet Perl module to assist with its internal functionality. We incorporated these modifications into MEME Suite version 4.11.0.

We characterize modified transcription factor binding sites using MEME-ChIP.^89^ It allows us to rapidly assess the main software outputs of interest: MEME^62^ and DREME,^90^ both for *de novo* motif elucidation;CentriMo,^55,91^ for the assessment of motif centrality; SpaMo,^92^ to assess spaced motifs (especially relevant for multipartite motifs);and Find Individual Motif Occurrences (FIMO).^93^

We mainly focus upon CentriMo^55^ for the analysis of our results. It permits inference of the direct DNA binding affinity of motifs, by assessing a motif’s local enrichment. In our case, we scan peak centres with PWMs, for the best match per region. We generate the PWMs used from MEME-ChIP, by loading the JASPAR 2014^94,95^ core vertebrates database, in addition to any elucidated *de novo* motifs from MEME or DREME. CentriMo counts and normalizes the number of sequences at each position of the central peaks to estimate probabilities of central enrichment. CentriMo smooths and plots these estimates, using a one-tailed binomial test to assess the significance of central enrichment.^55^

MEME-ChIP^89^ can yield repetitive motifs, without masking of low complexity sequences. Existing masking programs do not support modified genomes, and we accordingly mask the assembly, prior to modification with Cytomod. We use this masking only for downstream motif analyses. We use tandem repeat finder (TRF)^96^ (version 4.07b in mice and 4.09 in humans) to mask low complexity sequences. We used the following parameters: `2 5 5 80 10 30 200 -h -m -ngs`, from published TRF parameter optimizations.^97^ For version 4.09, to ensure compatibility with GRCh38/hg38 or larger future genomes, we increased the maximum “expected” tandem repeat length to 12 000 000, adding -l 12.

We ran MEME-ChIP,^89^ against Cytomod genome sequences for regions pertaining to ChIP-seq peaks from transcription factors of interest. For this analysis, we used the published protocol for the command-line analysis of ChIP-seq data.^98^ We employed positive controls, in two opposite directions, to assess the validity of our results. We use c-Myc as the positive control for an unmethylated binding preference.^25,26^ For this control, we used ChIP-seq data from a stringent streptavidin-based genome-wide approach with biotin-tagged Myc in mESCs from Krepelova et al.^58^ (GSM1171648). Also, we used murine erythroleukemia and CH12.LX Myc Mouse ENCODE samples (ENCFF001YJE and ENCFF001YHU). Conversely, we used both ZFP57 and C/EBPβ as positive controls for methylated binding preferences.^21–24^ For C/EBPβ, we used Mouse ENCODE ChIP-seq data, conducted upon C2C12 cells (ENCFF001XUT) or myocytes differentiated from those cells (ENCFF001XUR and ENCFF001XUS). Also, we used one replicate of ZFP57 peaks provided by Quenneville et al.^22^. When processing this replicate, we used the same parameters as for our other ZFP57 samples, except for employing default MACS stringency (*q* = 0.05). The reduced peak calling stringency allowed us to ensure sufficient peaks for this older, lower-coverage, dataset. We constructed a ZFP57 BED file using BEDTools^78^ (version 2.17.0) to subtract the control influenza hemagglutinin (HA) ChIP-seq (GSM773065) from the target (HA-tagged ZFP57: GSM773066).We retain only target regions with no overlap with any features implicated by the control file, yielding 11 231 of 22 031 features.

### Modified binding preferences of ZFP57

We used ZFP57 ChIP-seq data, provided by Strogantsev et al.^23^ (GSE55382), to examine the modified binding preferences of that transcription factor. Strogantsev et al.^23^ derived these 40 bp single-end reads from reciprocal F1 hybrid Cast/EiJ × C57BL/6J mESCs (BC8: sequenced C57BL/6J mother × Cast father and CB9: sequenced Cast mother × C57BL/6J father).

We re-processed the ZFP57 data to obtain results for NCBI m37/mm9. We performed this reprocessing similarly to some of the Mouse ENCODE datasets, to maximize consistency for future Mouse ENCODE analyses. We obtained raw FASTQs using Sequence Read Archive (SRA) Toolkit’s fastq-dump. Then, we aligned the FASTQs using Bowtie^79^ (version 1.1.0; `bowtie -v 2 -k 11 -m 10 -t --best --strata`). We sorted and indexed the BAM files using Sambamba.^76^ Finally, we called peaks, using the input as the negative enrichment set, using MACS 2^61^ (version 2.0.10) `callpeak`, with increased stringency (*q* = 0.00001), with parameters: `--qvalue 0.00001 --format ‘BAM’ --gsize ‘mm’`. This resulted in 90 478 BC8 and 56 142 CB9 peaks.

We used the ChIPQC^99^ Bioconductor^100^ package to assess the ChIP-seq data quality. We used the two control and two target runs for each of BC8 and CB9. Then, we used `ChIPQC(samples, consensus=TRUE, bCount=TRUE, summits=250, annotation=“mm9”, blacklist=“mm9-blacklist.bed.gz”, chromosomes = chromosomes)`. We set the utilized list of mouse chromosomes to only the canonical 19 autosomal and 2 sex chromosomes. Using a blacklist, we filtered out regions that appeared uniquely mappable, but empirically show artificially elevated signal in short-read functional genomics data. We obtained the blacklist file from the NCBI m37/mm9 ENCODE blacklist website (https://sites.google.com/site/anshulkundaje/projects/blacklists).^52^ The BC8 data had 13.7 % fraction of reads in peaks (FRiP) and the CB9 data had 9.12 % FRiP. Additionally, we performed peak calling at the default *q* = 0.05. This resulted in many more peaks for both BC8 (197 610 peaks; 27.6 % FRiP) and CB9 (360 932 peaks; 19.7 % FRiP). The CB9 sample had a smaller fraction of overlapping reads in blacklisted regions (RiBL). At the default peak calling stringency, BC8 had an RiBL of 29.7 %, while CB9 had only 8.38 %. This likely accounts for our improved results with the CB9 replicate (Figure S1).

Additionally, we analyzed three ZFP57 ChIP-seq replicates (100 bp paired-end reads) pertaining to mESCs in pure C57BL/6J mice.^101^ We paired each replicate with an identically-conducted ChIP-seq in a corresponding sample, which lacked ZFP57 expression (ZFP57-null controls).

For the pure C57BL/6J data, we used the same protocol as for the hybrid data, except for the following three differences. First, instead of input as the negative control, we used the ZFP57-null ChIP-seq data.We ran Bowtie in paired-end mode (using `-1` and `-2`). Second, we omitted the Bowtie arguments `--best --strata`, which do not work in paired-end mode. Instead, we added `-y --maxbts 800`, the latter of which we set with `--best`’s value, in lieu of the default threshold of 125. Third, we set MACS to paired-end mode (option `-f BAMPE`). This resulted in very few peaks, however, when processed with the same peak-calling stringency as the hybrid data (at most 1812 peaks) and FRiP values under 2 %. Even when we used the default stringency threshold, we obtained at most 4496 peaks, with

FRiP values of around 4.5 %. Nonetheless, we still observed the expected preference for methylated motifs (Figure S1).

### Processing of additional OCT4 and n-Myc datasets

We additionally used OCT4 and n-Myc ChIP-seq data, from Yin et al.^36^ (GSE94634). Except as indicated below, we processed these in the same manner as our ZFP57 data. For these transcription factors, we used mouse data for three conditions per factor. These conditions consist of a wildtype sample, a triple-knockout sample for *TET1+TET2 +TET3,* and a triple-knockout sample for *DNMT1+DNMT3A+DNMT3B.* OCT4 has two antibodies, both replicated, across all three conditions. Because of pooling of both replicates for one antibody in the TET triple-knockout condition, this leads to 3 conditions × 2 antibodies × 2 replicates −1 pooled replicate = 11 OCT4 samples. The n-Myc came from only a single antibody, resulting in 3 conditions × 2 replicates = 6 samples. We used the provided IgG samples as negative controls for peak calling for this dataset. As discussed previously,^36^ we also called peaks for matching samples against each of their mouse, rabbit, and goat IgG samples.

### Comparing motif modifications, using hypothesis testing

To directly compare various modifications of motifs to their cognate unmodified sequences, we adopted a hypothesis testing approach. One can derive motifs of interest from a *de novo* result that merits further investigation. Often, however, researchers identify motifs of interest using prior expectations of motif binding preferences in the literature, such as for c-Myc, ZFP57, and C/EBPβ. For every unmodified motif of interest, we can partially or fully change the base at a given motif position to some modified base (Table 3).

To directly compare modified hypotheses to their cognate unmodified sequences robustly, we tried to minimize as many confounds as possible. We fixed the CentriMo central region width (options `--minreg 99 --maxreg 100`). We also compensated for the substantial difference in the background frequencies of modified versus unmodified bases. Otherwise, vastly lower modified base frequencies can yield higher probability and sharper CentriMo peaks, since when CentriMo scans with its “log-odds” matrix, it computes scores for nucleobase *b* with background frequency f(*b*) as

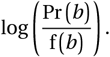

To compensate for this, we ensured that any motif pairs compared have the same length and similar relative entropies. To do this, we used a larger motif pseudocount for modified motifs (using CentriMo option `--motif-pseudo`).We computed the appropriate pseudocount, as described below, and provided it to iupac2meme. We set CentriMo’s pseudocount to 0, since we had already applied the appropriate pseudocount to the motif. We seek to normalize the average relative entropies of the PWM columns between two motifs.

The relative entropy (or Kullback-Leibler divergence), D_RE_, of a motif *m* of length |*m*|, with respect to a background model *b* over the alphabet *A*, of size |*A*|, is^102^

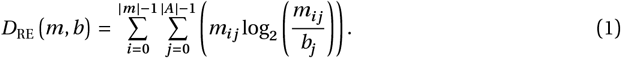

For each position, *i*, in the motif, the MEME Suite adds the pseudocount parameter, *α*, times the background frequency for a given base, *j*, at the position: 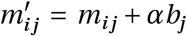. This omits the effective number of observed sites, which the MEME Suite also accounts for, essentially setting it to 1.

Accordingly, to equalize the relative entropies, we needed only substitute 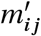 for each *m_ij_* in Equation 1 and then isolate α. In this process, we solve for *α*, by equating D_RE_ for the unmodified motif with that of the modified motif, substituting as above, while holding *α* for the unmodified motif constant. If we proceed in this fashion, however, our pseudocount would depend upon the motif frequency at each position and the background of each base in the motif. Instead, we can make a number of simplifying assumptions that apply in this particular case. First, the unmodified and modified motifs we compare differ only in the modified bases, which in this case, comprise only C or G nucleobases, with a motif frequency of 1. Additionally, we set the pseudocount of the unmodified motif to a constant 0.1 (CentriMo’s default). Thus, the pseudocount for a single modified base is the value α, obtained by solving, for provided modified base background frequency *b_m_* and unmodified base frequency *b_u_*:

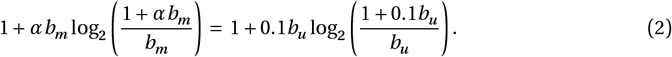

Equation 2 only accounts for a single modification, however, on a single strand. For complete modification, we also need to consider the potentially different background frequency of the modified bases’ complement. Thus for a single complete modification, with modified positions *m*_1_ and *m*_2_ and corresponding unmodified positions *u*_1_ and *u*_2_, modified base background frequencies *b*_*m*_1__, *b*_*m*_2__ and unmodified base frequencies *b*_*u*_1__, *b*_*u*_2__, we obtained

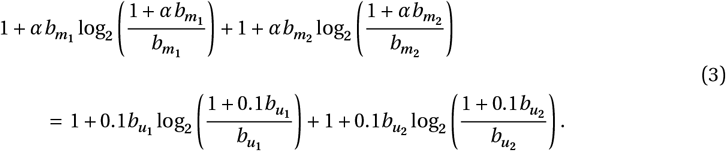

We numerically solved for *α* in Equation 3 for each modified hypothesis, using fsolve from SciPy.^103^ Finally, we may have multiple modified positions. We always either hemi-modify or completely modify all modified positions, so the pseudocount is the product of modified positions and the *α* value from Equation 3.

The pseudocount obtained in this fashion does not exactly equalize the two motif’s relative entropies, since we do not account for the effect that the altered pseudocount has upon all the other positions of the motif. It does, however, exactly equalize the relative entropies per column (RE/col, as defined by Bailey et al.^102^) of the modified versus unmodified motifs, which suffices to ensure correctly normalized comparisons.

Then, we performed hypothesis testing for an unmodified motif and all possible 5mC/5hmC modifications of all CpGs for known modification-sensitive motifs for c-Myc, ZFP57, and C/EBPβ. These modifications consist of the six possible combinations for methylation and hydroxymethylation at a CpG, where a CpG is not both hemi-methylated and hemi-hydroxymethylated. These six combinations are: `mG, C1, m1, hG, C2`, and `h2`. For c-Myc, we constructed modified hypotheses from the standard unmodified E-box: `CACGTG`. For ZFP57, we tested the known binding motif, as both a hexamer (`TGCCGC`) and as extended heptamers (`TGCCGCR` and `TGCCGCG`).^22,23^ We additionally tested motifs that occurred frequently in our *de novo* analyses, `C(C/A)TGm1(C/T)(A)`. We encoded this motif as the hexamer `MTGCGY` and heptamers, with one additional base for each side: `CMTGCGY` and `MTGCGYA`. This encoding permitted direct comparisons to the other known ZFP57-binding motifs of the same length. Finally, for C/EBPβ we tested the modifications of two octamers: its known binding motif (TTGCGCAA) and the chimeric C/EBP|CRE motif (TTGCGTCA).^21^

Using CentriMo, we assessed motifs for their centrality within their respective ChIP-seq datasets. Then, we computed the ratio of CentriMo central enrichment p-values, adjusted for multiple testing,^55^ for each modified/unmodified motif pair. For numerical precision, we computed this ratio as the difference of their log values returned by CentriMo. This determines if the motif prefers a modified (positive) or unmodified (negative) binding site.

We conducted hypothesis testing across all four replicates of mouse WGBS and oxWGBS data, for a grid search of modified base calling thresholds. The levels output by MLML,^83^ allowed us to obtain these thresholds. We interpret these values as our degree of confidence for a modification occurring at a given locus. We conducted our grid search from 0.01–0.99 inclusive, at 0.01 increments. Finally, we plotted the ratio of CentriMo p-values across the different thresholds, using Python libraries Seaborn^104^ and Pandas.^105^ Then we extended this combinatorial hypothesis testing approach, across all JASPAR and *de novo* motifs from our Mouse and Human ENCODE datasets, at our 0.3 and 0.7 selected thresholds.

### Assessment of transcription factor familial preferences

We used TFClass^56,57^ (downloaded November 8, 2017) to categorize and group analyzed transcription factors into families and super-families. We used Pronto^106^ (version 0.2.1) to parse the TFClass ontology files.

For transcription factors either not categorized at time of analysis or that yielded inexact matches, we manually assigned them to families and super-families. We performed curation by searching one or more of GeneCards,^107^ Genenames.org,^108^ UniProt,^109^ Gene3D,^110^ InterPro,^111^ Pfam,^112^ SMART,^113^ and SUPERFAMILY.^114^

We manually re-named a number of transcription factors in the family assignment to match the names used elsewhere in our data, largely removing hyphens for consistency, creating a group for POLR2A, and adding a number of missing transcription factors. The following factors (with asterisks denoting any suffix and slashes denoting synonymous factors) underwent this manual annotation: RAD21, REC8, SCC1, ZC3H11*, CHD*, NELFE, PAH2, SIN3*, PIAS*, ZMIZ*, KLHL, HCFC*, EP300, TCF12, TIF1*/TRIM24/TRIM28/TRIM33, SMC*, KAT2A/GCN5, and SMARCA4/BRG1.

We aggregated all hypothesis testing results across either mouse or K562 datasets distinctly. Grouped by modification type (m or h), we aggregated across stringency (0.3 or 0.7), replicate of origin, and unmodified hypothesis. When comparing a modified hypothesis pair to its unmodified counterpart, different replicates of data may produce different scores. In this instance, to aggregate multiple similar hypothesis tests, we took the maximum absolute value score. For each transcription factor, we retained only the most statistically significant (“top-1”) or top three most statistically significant (“top-3”) hypothesis pairs. We omitted hypothesis pairs that lack statistical significance (p-value ≥ 0.05).

### Motif clustering of modified binding preferences

We used Regulatory Sequence Analysis Tools (RSAT) `matrix-clustering`^115^ to hierarchically cluster similar motifs. For each transcription factor, we clustered each of its unmodified motifs, alongside their modified counterparts. These motifs matched the set of hypothesis pairs for that factor.

We partitioned each transcription factor’s motifs into unmodified-preferring (score < –*ε*), modified-preferring (score > *ε*), and those without any substantive preference (-*ε* ≤ 0 ≤ *ε*). Here, we set ε = 5, to ignore any near-neutral preferences.

After this, we removed duplicate hypothesis pairs, selecting only those with the scores furthest away from zero. Then, we plotted these clusters, annotated by their score, in a treemap^116^ plot. We created this plot using R^117^ (version 3.5.1) ggplot2’s^118^ treemapify^119^ and Python (version 2.7.15) Pandas (version 0.22.0)^105^ data structures through rpy2^120^ (version 2.8.6).

We designed a colour scheme to highlight motifs with strong preferences. To do this, we used white for scores of 80, above or below 0 (namely, from −80–80). This represented an expansion of disregarded motif from the *ε* = 5 threshold used above. We also kept the colour ramp linear within a score of 20 on either side of 0. Outside of this range around the centre, the ramp becomes logarithmic.

To further highlight the rarer and lower scores occurring for motifs preferring to bind in modified contexts, we altered the mid-point of the colour scheme. To do this, we re-centred the colour scheme, shifting it −10, thereby biasing it towards modified contexts, in shades of red. The re-centring offset skews the entire colour scheme toward red hues, including moving the white regions accordingly.

### Validation of our OCT4 findings, using CUT&RUN

After finding that OCT4 had a number of both methyl- and hydroxymethyl-preferring motifs, we performed CUT&RUN^49,50^ on mESCs, targeting OCT4. We also performed CUT&RUN for a matched IgG, for use as a background during peak calling. We performed CUT&RUN on mESCs, targeting OCT4. We subjected the resultant DNA to 3 workflows: conventional library preparation for sequencing on an Illumina platform, bisulfite sequencing, and Nano-hmC-Seal-seq.^121^ Using a NovaSeq 6000 (Illumina), we sequenced the resulting libraries from the 3 workflows, using a paired-end 2 × 150 bp read configuration (Princess Margaret Genomics Centre, Toronto, ON, Canada).

#### Cell lines

Using feeder-free conditions, we grew male E14 murine embryonic stem cells^122^ on 10cm plates gelatinized with 0.2 % porcine skin gelatin type A (Sigma, St. Louis, MO, USA) at 37 °C and 5 % CO_2_. We cultured cells in N2B27+2i media.^123^ Briefly, this media contains DMEM/F12^124^ (Sigma) and Neurobasal media (ThermoFisher, Waltham, MA, USA), supplemented with 0.5× B27 (Invitrogen, Waltham, MA, USA), 1× N-2 Supplement, 50μmol/l 2-mercaptoethanol (ThermoFisher), 2 mmol/l glutamine (ThermoFisher), Leukemia Inhibitory Factor (LIF), 3 μmol/l CHIR99021 glycogen synthase kinase (GSK) inhibitor (p212121, Boston, MA, USA), and 1 μmol/l PD0325091 mitogen-activated protein kinase/extracellular signal-regulated kinase kinase (MEK) inhibitor (p212121). We passaged cells every 48 h using trypsin (Gibco, Waltham, MA, USA) and split them at a ratio of ≈ 1: 8 with fresh medium. We conducted routine anti-mycoplasma cleaning (LookOut DNA Erase spray, Sigma) and screened cell lines by PCR to confirm no mycoplasma presence.

#### CUT&RUN assay

We performed CUT&RUN as described elsewhere^125–127^ using recombinant Protein A-micrococcal nuclease (pA-MN). Briefly, we extracted nuclei from ≈ 4500000 embryonic stem cells using a nuclear extraction buffer comprised of 20 mmol/l 4-(2-hydroxyethyl)-1-piperazineethanesulfonic acid (HEPES)-KOH,^128^ pH 7.9, 10mmol/l KCl, 0.5 mmol/l spermidine, 0.1 % Triton X-100, 20 % glycerol, and freshly added protease inhibitors. We bound the nuclei to 500 μl pre-washed lectin-coated concanavalin A magnetic beads (Polysciences, Warrington, PA, USA). Then, we washed beads in binding buffer (20mmol/l HEPES-KOH, pH 7.9, 10 mmol/l KCl, 1 mmol/l CaCl_2_ MnCl_2_). We pre-blocked immobilized nuclei with blocking buffer (20 mmol/l HEPES, pH 7.5, 150 mmol/l NaCl, 0.5 mmol/l spermidine, 0.1 % bovine serum albumin (BSA), 2 mmol/l EDTA, fresh protease inhibitors). We washed the nuclei once in wash buffer (20 mmol/l HEPES, pH 7.5, 150mmol/l NaCl, 0.5 mmol/l spermidine, 0.1 % BSA, fresh protease inhibitors). Following this, we incubated nuclei in wash buffer containing primary antibody (anti-Oct4, Diagenode (Denville, NJ, USA) cat. no. C15410305 (lot A2089P) or anti-IgG, Abcam (UK) cat. no. ab37415; RRID: AB_2631996) for 1 h at 4 °C with rotation. Then, we incubated in wash buffer containing recombinant pA-MN for 30 min at 4 °C with rotation.

Using an ice-water bath, we equilibrated samples to 0°C and added 3 mmol/l CaCl_2_ to activate pA-MN cleavage. Then, we performed sub-optimal digestion, at 0°C for 30min. As described in Step 31 of Skene et al.^50^, we intentionally conducted digestion at a temperature lower than optimal, to prevent otherwise unacceptable background cleavage levels.^49,50^ We chelated digestion with 2XSTOP+ buffer (200mmol/l NaCl, 20mmol/l EDTA, 4mmol/l ethylene glycol-bis(β-aminoethyl ether)-*N,N,N,’*-tetraacetic acid (EGTA), 50μg/ml RNase A, 40μg/ml glycogen, and 1.5pg MNase-digested *Saccharomyces cerevisiae* mononucleosome-sized DNA spike-in control).

After RNase A treatment and centrifugation, we released and separated genomic fragments. We digested protein using proteinase K. Finally we purified DNA using phenol:chloroform:isoamyl alcohol extraction, followed by ethanol precipitation.

#### Library preparation for bisulfite sequencing

We prepared our bisulfite sequencing library using 30 ng of CUT&RUN DNA. We used the Ultra II Library Preparation Kit (New England Biolabs (Canada) (NEB), cat. no. E7645L) following manufacturer’s protocol, with some modifications. In brief, after end-repair and A-tailing, we ligated NEBNext methylated adapters for Illumina (NEB, cat. no. E7535) at a final concentration of 0.04μmol/l onto the DNA, followed by incubation at 20°C for 20 min. Post adapter incubation, we subjected adapters to USER enzyme digestion at 37°C for 15 min prior to clean up using AMPure XP Beads (Beckman Coulter, cat. no. A63881).

We bisulfite-converted adapter-ligated CUT&RUN DNA using Zymo Research EZ DNA Methylation Kit (Zymo, Irving, CA, USA, cat. no. D5001) following the alternative protocol for the Infinium Methylation Assay (Illumina). Briefly, we added 5 μl of M-Dilution buffer to purified adapter-ligated DNA, and adjusted total sample volume to 50μl with sterile molecular grade water. We incubated samples at 37°C for 15min, prior to the addition of 100μl of CT Conversion Reagent. We further incubated samples, prior to purification, at (95 °C for 30 s, 50°C for 60 min) for 16 cycles, then at 4 °C for at least 10 min, following the manufacturer’s protocol. Finally, we eluted samples in 23 μl molecular grade water.

We amplified the bisulfite-converted DNA using 2× HiFi HotStart Uracil+ ReadyMix (KAPA, Wilmington, MA, USA, cat. no. KK2801), and unique dual index primers (NEB, cat. no. E6440S) in a final volume of 50μl. We performed this using the following PCR program: 98°C for 45 s, followed by 17 cycles of: 98°C for 15s, 65°C for 30s, 72°C for 30s, and final extension at 72°C for 60s. We purified and dual size selected amplified libraries using AMPure (Beckman Coulter, ON, Canada) XP Beads at 0.6× to 1.0× bead ratio, eluted in a volume of 20μl.

#### Library preparation for hmC-Seal sequencing

We performed library preparation using the NEB Ultra II Library Preparation Kit (NEB, cat. no. E7645L) on 30 ng CUT&RUN DNA, as per the manufacturer’s protocol, with the below modifications. In brief, after end-repair and A-tailing, we purified adapter ligated DNA using AMPure XP Beads at a 0.9× ratio and eluted in 11.5 μl sterile water.

We added three spike-in DNA controls to the adapter ligated DNA to assess specific enrichment of modified DNA fragments. Controls consisted of 0.2 ng/μl working stocks of unmethylated and methylated *Arabidopsis* DNA spike-in controls from the Diagenode DNA methylation control package (cat. no. C02040012). They also included the 5hmC spike-in control DNA (amplified from the *APC* promoter) from the Active Motif Methylated DNA standard kit (Active Motif, cat. no. 55008). We combined 0.3 ng of each spike-in DNA in a final volume of 4.5 μl per experimental sample. We mixed the adapter ligated DNA with the spike-in DNA mix. Then we aliquoted 1.6 μl of this mix into a separate PCR tube and stored it at −20°C, as an input control.

We 5hmC-glucosylated the remaining 14.4 μl CUT&RUN DNA mixed with spike-in controls, as previously described,^121^ with the below modifications. Briefly, we 1: 1 diluted a 3μmol/l stock of uracil diphosphate (UDP)-azide-glucose (Active Motif, cat. no. 55020) in 1× phosphate-buffered saline (PBS) to establish a working stock of 1.5 μmol/l for Mastermix preparation. We prepared a 20 μl glucosylation Mastermix per experimental sample consisting of the following: 14.4 μl CUT&RUN DNA mixed with spike-in controls, 50μmol/l HEPES (pH 8.0), 25 mmol/l MgCl_2_, 0.1 mmol/l UDP-azide-glucose and 1 U of T4 Phage β-glucosyltransferase (NEB, cat. no. M0357L). We incubated the mix for 1 h at 37°C, to promote glucosylation.

Then, we performed biotinylation of azide-labelled 5hmC residues of the glucosylated DNA fragments. In sterile water, we prepared 20mmol/l dibenzocyclooctyne-PEG4-biotin conjugate (Bioscience, cat. no. CLK-A105P4-10) and stored it in one-time use aliquots at −20°C, to avoid freeze-thaw. We mixed 20μl of glucosylated DNA with 1.8 μmol/l dibenzocyclooctyne-PEG4-biotin in a final reaction volume of 22 μl, then incubated 2 h at 37°C to promote biotinylation. Then, we prepared MicroSpin P-30 Gel Columns (Bio-Rad, Hercules, CA, USA, cat. no. 7326223), following the manufacturer’s protocol, and used them to purify total DNA fragments from reaction components. Briefly, we loaded the sample onto the column, then centrifuged 4 min at 1000 × g to elute purified DNA sample in 22 μl of Tris buffer.

To specifically capture biotinylated 5hmC DNA fragments, we prepared 2× binding and washing (B&W) buffer (10 mmol/l Tris-HCl, 10 mmol/l EDTA, 2 mol/l NaCl). Using 20 μg of MyOne Streptavidin C1 Dynabeads (ThermoFisher, cat. no. 65001), we re-suspended in 0.2 ml of 1× B&W buffer per experimental sample to wash beads. We subjected beads to 3 total washes, then re-suspended to a final volume of 22 μl per sample in 2× B&W buffer. We added 22 μl of purified total DNA fragments to 22 μl of washed beads, then incubated 15 min under gentle rotation to promote streptavidin-biotin binding.

To isolate beads containing streptavidin-bound biotinylated DNA fragments, we incubated them on magnet for 3 min. Then, we washed the beads 3 times with 1× B&W buffer to remove non-biotinylated DNA fragments lacking 5hmC. Finally, we re-suspended the beads in 50μl of low TE buffer.

We conducted quantitative PCR (qPCR) to compare enrichment of 5hmC spike-in after biotin enrichment relative to input sample stored earlier. qPCR in the form of a 10μl reaction, consisted of 1× SYBR Fast qPCR Mastermix (KAPA, cat. no. KK4601), 1 μl of template DNA, and primers at a final concentration of 0.3μmol/l. We set up different reactions for each primer set to detect each spike-in control DNA separately. We used template-specific forward and reverse primers. For 5hmC, we quantified spike-in DNA fragment from the Active Motif Methylated DNA standard kit. We also quantified the methylated or unmethylated *Arabidopsis* DNA spike-in controls from the Diagenode DNA Methylation Control package kit. We amplified with the following PCR program: 98°C for 30 s, followed by 40 cycles of 98 °C for 15 s and 60 °C for 15 s (with image capture), ending with melt curve analysis.

To generate bead-free template for library DNA amplification, we established a PCR reaction mix containing 0.3 μmol/l of unique dual index primers (NEB, cat. no. E6440S), 1× NEBUltra II Q5 MM, and DNA/bead template for a final volume of 100μl per sample. We split samples into 2× 50μl reactions and amplified using the following PCR program: 98 °C for 30 s, followed by 5 cycles of 98 °C for 10 s, 60 °C for 75 s, ending with a hold at 4 °C. Then, we transferred reaction tubes to a magnetic rack and transferred bead-free supernatant to new PCR tubes. To amplify DNA libraries for a maximum of 16 cycles total (including initial 5 cycles), we used the same PCR conditions for the bead-free template. Then, we dual size selected DNA libraries using AMPure XP beads at 0.7× to 1.0× ratio, as described in the library preparation for bisulfite sequencing.

#### Sample sequencing

We performed library preparation of 5 ng of CUT&RUN DNA, following the NEB Ultra II Library Preparation Kit (cat. no. E7645L) manufacturer’s protocol. We used different NEB dual indices for each sample (Table 4). We sequenced all libraries on a NovaSeq 6000 sequencing system using a SP flow cell run in standard mode, with paired-end 2 × 150 bp read length configuration. This allowed us to obtain the desired number of reads per sample (Table 4).

**Table 4.** CUT&RUN samples used in our experiments, with their sequencing technique, indices, and target read details. Target reads represent the number of single-end equivalent Illumina passing-filter read estimates we sought to obtain.

| Sample name | Technique | i7 Index | i5 Index | Target reads<br>(millions of<br>single reads) |
| --- | --- | --- | --- | --- |
| OCT-4_Pool_1 | CUT&RUN | TCTAGGAG | AGGTCACT | 40 |
| OCT-4_Pool_2 | CUT&RUN | TGCGTAAC | GATAGGCT | 40 |
| IgG_Pool_1 | CUT&RUN | CTTGCTAG | GGAGATGA | 20 |
| OCT-4_Pool_1_BS | CUT&RUN-BS | AGCGAGAT | GATACTGG | 180 |
| OCT-4_Pool_2_BS | CUT&RUN-BS | TATGGCAC | TCTCGCAA | 180 |
| IgG_Pool_1_BS | CUT&RUN-BS | GAATCACC | CTTCGTTC | 40 |
| Oct-4_hmeseal_rep_1 | CUT&RUN-5hmC | GTAAGGTG | GCAATTCG | 150 |
| Oct-4_hmeseal_rep_2 | CUT&RUN-5hmC | CGAGAGAA | TCTCTTCC | 150 |

#### Data processing

We performed base calls using Real-Time Analysis (RTA) (version 3.4.4). Using bcl2fastq (version 2.20), we converted Binary Base Call (BCL) files to FASTQ files.

We processed the CUT&RUN sequences as follows. Before alignment, we trimmed adapter sequences with fastp (version 0.19.4).^129^ We assessed sequencing data quality using FastQC (version 0.11.8),^130^ Picard^131^ (version 2.6.0) `CollectInsertSizeMetrics`, QualiMap^132^ (version 2.2) `bamqc`, Preseq^133^ (version 2.0.0) `bound_pop` and `lc_extrap`, DeepTool^s134^ (version 3.1.3), and MultiQC^135^ (version 1.7). For tools requiring Java, we used Java SE 8 Update 45. For tools requiring, Python we used version 2.7.12, except as otherwise noted.

We aligned reads to GRCm38/mm10 with Bismark^73^ (version 0.22.3; for 5mC or 5hmC sequences). Bismark used Bowtie 2^79–81^ (version 2.4.1; and for conventional sequences), SAMtools^82^ (version 1.10), and BEDTools^78^ (version 2.29.2). We used Bismark’s default parameters, save those controlling output destinations and use of multiple cores, and parameters passed to Bowtie 2, as described below.

We used Bowtie 2 parameters as recommended,^136^ excepting increasing alignment sensitivity, and specifying implied or default parameters. Therefore, we used the parameters `-D 20 -R -N 1 -L 18 -i S,1,0.25` for increased sensitivity, slightly more so than the `--very-sensitive-local` preset. We used `-I 10` for a minimum fragment length of 10 bp and `-X 700` for a maximum fragment length of 700 bp, as recommended.^50,136^ This range of fragment lengths included those we selected for during library preparation (30 bp–280 bp).We also used the parameters `--local --phred33 --no-unal --no-discordant --no-mixed`. For alignments used for calculating the spike-in coefficient, we did not permit dovetailing (`--no-dovetail`) nor overlaps (`--no-overlap`), as recommended.^50^

For post-processing, we used Sambamba^76^ (version 0.7.1), including marking duplicates. Where applicable, we performed spike-in calibration as described by Meers et al.^136^.

For our final OCT4 results, we did not use our *Saccharomyces cerevisiae* spike-in calibrated data. In the unmodified context, the spike-in calibrated data made little difference. In the modified context, insufficient modified bases in the spike-in probably prevented us from properly calibrating.

We called peak summits using MACS 2^61^ (version 2.1.2). We ensured that the input only included reads with insert sizes ≤120 bp, as recommended,^50,136^ by using DeepTools^134^ (version 3.1.3) `alignmentSieve`.

For data not calibrated with spike-in, we used MACS 2 `callpeak`, specifying treatment and control inputs and outputs as usual. We used the additional MACS 2 parameters, `--buffer-size 1000000 --format ‘BAMPE’ --gsize ‘mm’ --qvalue 0.05 --call-summits --bdg --SPMR`.

For spike-in calibrated data, we used advanced MACS sub-commands, constructed to yield a peak calling scheme that worked well for CUT&RUN datasets. Specifically, we used pileup on BAMPE input, then bdgopt to multiply by the scaling factor defined by the spike-in calibration. Then, we added a pseudocount of 1.0 to mimic the default workflow, using bdgcmp, specifying `--pseudocount ‘0.0’ --method ‘qpois’`, followed by `bdgpeakcall, with --cutoff` -ln(0.05)/ln(10). This cutoff parameter represented the usual q-value cutoff of 0.05 converted to - log_10_ space.

For bisulfite-converted data, we extracted and called peaks only upon methylated reads (filtering through Sambamba using Bismark’s added “XM:Z:” tag). We regarded all hmC-Seal-seq reads as completely hydroxymethylated.

Finally, we used MEME-ChIP (version 4.11.2.1, with Perl version 5.18.1),^89^ with DREME,^90^ as previously described, on a TRF-masked genome (same parameters as before, using version 4.9). For this particular processing, we used SAMtools^82^ version 1.3.1 and BEDTools^78^ version 2.27.1.

## Discussion

We developed a method of creating modified genomic sequences at suitable thresholds, using our tool Cytomod. We have added expanded alphabet capabilities to the widely-used MEME Suite,^48^ a set of software tools for the sequence-based analysis of motifs. This included extending several of its core tools, including: MEME,^62^ DREME,^90^ and CentriMo,^55^ used in a unified pipeline through MEME-ChIP.^89^ We undertook further extension of all downstream analysis tools and pipelines, and most of the MEME Suite^48^ now supports arbitrary alphabets. Our approach has yielded a much greater understanding of transcription factor’s affinities and motifs in a modified genomic context. We validated our novel OCT4 binding site predictions, generating new high-quality binding site data, in both unmodified and modified genomic contexts.

We devised a hypothesis testing approach to enable more accurate comparisons between unmodified and modified motifs. Hypothesis testing, with equal central region widths and relative entropies, leads to more interpretable results than the standard CentriMo analyses, in that it permits a direct comparison of centrality p-values. These p-values help assess the statistical significance of the motif within the central region of its detected binding enrichment—a strong indicator of direct DNA binding.^55^ We often observed the expected outcomes for many replicates of conventional CentriMo runs with *de novo* motifs, such as with C/EBPβ (Figure 3 and Figure 4)and ZFP57 (Figure S4). We encountered instances, however, in which *de novo* CentriMo analyses did not show the expected motif binding preference. This occurred for c-Myc and for a small subset of ZFP57 CentriMo results pertaining to *de novo* motifs, despite the hypothesis testing robustly corroborating its expected preference for unmethylated DNA (Figure S1). Overall, our hypothesis testing framework allows for a more accurate comparison than a direct assessment of *de novo* motifs, which would be less well-controlled for technical biases.

Various biochemical complexities increase the difficulty of mapping cytosine modifications. These complexities include strand biases,^9^ populations of cells with different modifications at the same locus, and hemi-methylation.^137^ Our use of MLML^83^ to provide consistent estimators of modification, our relative entropy normalization, and our controlled hypothesis testing approach all help to minimize the impact of these challenges.

Cytosine modifications occur most frequently at CpG dinucleotides. Nevertheless, non-CpG 5mC nucleobases still exist, particularly in mESCs.^138,139^ Within a population of cells, at a given locus, unmodified nucleobases and different kinds of modified nucleobases often co-occur.^9^

Our methods ensure the comprehensive analysis of non-CpG modifications. While this can result in some modified hypotheses being unlikely to occur in some cell types, we can still evaluate and score those hypotheses in an unbiased and tissue-specific manner. Therefore, some motifs shown may be unlikely to occur, but will usually tend to have scores near 0. One can interpret such scores as a weak preference, should that motif be present. Our DNAmod database catalogues these and many other DNA modifications.^140^

We suspect that the inability of *de novo* analyses to reveal modified binding preferences primarily arises from being unable to integrate modified and unmodified motifs. Our *de novo* analyses cannot compensate for the large differences in modified versus unmodified background frequencies. *De novo* analysis involves some form of optimization or heuristic selection of sites—an inherently variable process. Modified motifs have particular characteristics that differ from most unmodified motifs. Most notably, they necessarily differ from the overall and likely local sequence backgrounds, as a result of the low frequency of modifications. Conversely, an unmodified genome sequence has a comparably uniform nucleobase background, and unmodified motifs usually appear within local sequence of highly similar properties to the motifs themselves.^141^ Accordingly, without specifically accounting for these confounds, modified motifs can get lost within a background of irrelevant unmodified motifs or one might not find comparable sets of motifs. Also, modified motifs that a *de novo* analysis finds might not be comparable to any unmodified counterpart. This may arise from the potential pairs of motifs having substantially different lengths, often with the modified motifs having significantly shorter length. Comparing motifs having sequence properties that often indicate a poor-quality motif also remains difficult. These properties include repetitious motifs, or off-target motifs, such as zingers—common contaminant motifs similar to CTCF, ETS, JUN, and THAP11.^142^ Hypothesis testing, with relative entropy normalization, can mitigate these concerns. Possibly, however, we often simply observe bona fide, but non-canonical, motifs. Non-canonical binding sites have far more abundance and importance than generally appreciated.^143^ Therefore, while our approach can yield biologically-relevant modified *de novo* motifs, one should not rely solely on these motifs’ binding preferences to conclusively establish a factor’s preference for modified DNA. Our hypothesis testing approach, however, helps mitigate the above biases, allowing for more robust comparisons.

Our method is robust in the face of parameter perturbations. There exists an inherent trade-off between a lower and higher modification calling threshold. The low threshold may yield more modified loci but potentially introduce false positives, while a higher threshold may prove too stringent to detect modified base binding preferences. Nonetheless, expanded-alphabet motif analysis across a broad range of modified base calling thresholds consistently led to the same expected results, across three transcription factors and a number of ChIP-seq and bisulfite sequencing replicates (Figure S1). We selected a lower threshold of 0.3, based primarily on the observation of increased variance and decreased apparent preference for unmethylated DNA for c-Myc below this threshold, across multiple replicates (Figure S1). We also selected an upper threshold of 0.7, based primarily on the rapid decrease in relative affinity for methylated over unmethylated motifs in ZFP57 (Figure 2) and, to a lesser extent, C/EBPβ (Figure S3). Furthermore, modification of peak calling stringency for a set of ZFP57 datasets did not negatively impact our ability to detect the affinity of ZFP57 for methylated DNA (Figure S2).

The consistency of our controls (c-Myc, ZFP57, and C/EBPβ) provides confidence in the ability of our method to detect and accurately characterize the effect of modified DNA on transcription factor binding. We applied our method to a diverse array of ChIP-seq data in order to identify biologically-meaningful binding preferences. We were also able to confirm OCT4 binding preferences, by generating new experimental data.

We found that motifs often enriched for hemi-modified, as opposed to completely-modified binding sites. These hemi-modified motifs often had more central enrichment, as measured by CentriMo, than those with complete modification of a central CpG dinucleotide (Figure 3; Figure 4). This appears surprising, because *in vitro* experiments have demonstrated that each modification usually has an additive effect for transcription factors that prefer modified DNA, resulting in completely modified motifs having the greatest affinity.^21,24^ This might imply that the hemi-methylated motifs arise from technical artifacts, either in the bisulfite sequencing data or from the methods used. Alternatively, the hemi-methylation events we detect may arise from asymmetric binding affinities of transcription factors for 5mC (and 5hmC). ZFP57, for example, has known asymmetric recognition of 5mC, with negative strand methylation more important than positive strand methylation in the TGCC**G**C motif.^24^ In addition, there exists evidence for a preference for hemi-methylation of the C/EBP half-site |`GmAA`.^59^ Therefore, the hemi-methylated motifs observed in some of our analyses, especially for C/EBPβ, may represent bona fide preferences. This would accord with similar findings in an independent analysis.^42^ Nevertheless, further work is needed to determine whether the hemi-methylated motifs we discover reflect an actual biological preference.

There exist few high-quality single-base resolution datasets of 5hmC, 5fC, and 5caC. We had previously attempted analyses using DNA modification data that did not have single-base resolution, such as from assays like methylation DNA immunoprecipitation (MeDIP),^144^ that did not employ single-base resolution methods.^145^ A lack of single-base resolution makes it difficult to create a discrete genome sequence with a reasonable abundance of the modification under consideration without biasing the sequence. This makes downstream analyses of transcription factor binding uninformative. Therefore, it is essential to have single-base resolution data, for any modifications that one wishes to analyze. Additionally, many single-base resolution datasets use some form of reduced representation approach that enriches CpGs, because this allows sequencing at reduced depth, while still capturing many DNA modifications. The use of reduced representation bisulfite sequencing data can lead to confounding factors, due to the non-uniform distribution of methylated sites surveyed. Accordingly, we recommend avoiding similar enrichment approaches for use with our framework.

The ChIP-seq data we used were not generated in the same cell type as the WGBS and oxWGBS data. While cell-type specificity might cause confounding effects, we consistently observed the expected preferences in transcription factor binding for the expected modification affinities across multiple ChIP-seq replicates, often in different cell types. Therefore, we expect that using ChIP-seq and WGBS data from different cell types will lead to meaningful results.

Although we predominantly the observed expected transcription factor binding preferences, in some limited instances we did not. For example, we found that USF1 and USF2 appear to have a subset of 5mC- and 5hmC-preferring motifs (Figure 5C). This contradicts previous *in vitro* work,^146^ which showed that USF1 prefers to bind neither 5mC nor 5hmC. Additionally, the same study suggested that while TCF3, a transcription factor related to USF1 and USF2, can undergo a conformational shift to bind to 5hmC, USF1 cannot. The *in vitro* work largely derived from structural preferences, however: it “only focused on the most obvious, steric and hydrogen bonding effects … [more] complex methods are required to explain [their] subtler [protein binding microarray] results”.^146^ Nonetheless, in their data, some of the C versus 5hmC and 5mC versus 5hmC z-score plots depict a few significant motif pairs with near equal preference for versus against hydroxymethylation. This indicates that their more general conclusion may not accurately sum up all of their data either; our results are more in accord than it would appear at first glance.

Our results for USF1 only had a single weakly positive-scoring motif containing a hydroxymethylated base: `AAAhYAmA`. This motif had only a slight preference to bind over its unmodified counterpart. This preference may instead have arisen primarily from the methylated base near the end of the motif. The low score might indicate that there is no strong preference; it could also suggest a technical artifact. All our other modified-preferring motifs for USF1 preferred to bind in a methylated context, with most showing only weak preferences. This stood in contrast to the many methylated motifs that showed strong preferences for their unmodified counterparts. Overall, these results suggest that not all USF1 motifs tend to bind in unmodified contexts, even if most of them may do so. USF2, however, has a non-negligible number of motifs that appear able to bind in a hydroxymethylated context, having 5hmC motifs that scored above 0.

Our limited assessment of transcription factor preferences across different families did not yield clear conclusions. Despite previous findings for specific families, like basic helix-loop-helix factors tending to prefer to bind to unmodified DNA,^27–31^ most conclusions in this area are ambiguous.^40–42,147^ One reason for this is the different motif groups that prefer unmethylated versus methylated binding for many transcription factors. This tends to confound binary categorization even for individual transcription factors. Across a whole family of transcription factors, making this binary call becomes even more difficult. Even closely related transcription factors, like USF1/2, often have different degrees of preferences and variable motifs. A second reason is observation bias with regards to transcription factors for which one can find binding data. There is a particular depletion of binding data for the large number of zinc finger transcription factors. This bias may account, at least partially, for the often observed greater number of unmodified-preferring factors.^20,36^ For an unbiased assessment of transcription factors across families, we need data on a less biased set of transcription factors.^36,148,149^

The MEME Suite’s^48^ new custom alphabet capability permits further downstream analyses of modified motifs. Our custom alphabet is provided together with this software, and is available both from the MEME Suite webpage and as the standalone MEME::Alphabet package. For example, one can find individual motif occurrences with FIMO^93^ or conduct pathway analyses with Gene Ontology for MOtifs (GOMO).^150^ Alternatively, one can use FIMO results for pathway analyses through tools like Genomic Regions Enrichment of Annotations Tool (GREAT)^151^ or biological enrichment of hidden sequence targets (BEHST).^152^ For further interpretation of the results, one can use downstream pathway analysis tools, such as Enrichment Map.^153,154^ This permits inference of implicated genomic regions and biological pathways.

We designed all of our software so that others can readily extend our approach to additional DNA modifications. Technology now allows the detection of a number of DNA modifications^140^ at high resolution, such as 5-hydroxymethyluracil (5hmU), 5-formyluracil (5fU), 8-oxoguanine (8-oxoG), and 6-methyladenine (6mA), many of which occur in diverse organisms.^155–157^ We provide recommendations in Appendix A for the nomenclature of these modified nucleobases, among others. We used these recommendations in our database of DNA modifications, DNAmod.^140^ The Global Alliance for Genomics and Health (GA4GH)^158^ has also adopted these recommendations for use in sequence alignment/map (SAM) and BAM formats.^82^

It will be important to characterize modified binding affinities *in vivo,* in addition to the more abundant *in vitro* approaches, such as high-throughput systematic evolution of ligands by exponential enrichment (HT-SELEX) and DNA affinity purification sequencing (DAP-seq).^34^ While the *in vitro* approaches contribute to our improved understanding of the underlying biophysics, only *in vivo* analyses can directly assess the actual cellular binding events that lead to differences in gene expression pattern. By using available ChIP-seq data, our work contributes to this effort and bolsters it by providing a transcription factor CUT&RUN dataset that directly assesses unmodified and modified binding states. These data represent a unique type of experiment, one needed to fully understand the role of methylation and hydroxymethylation in transcription factor binding.

We provide a framework for transcription factor binding motif analyses on sequences containing DNA modifications. Our approach’s ability to reproduce known transcription factor binding affinities and the validation of our predictions for OCT4 suggest that these methods meaningfully predict the modification sensitivity of transcription factors. One can use our approach to analyze a wide array of transcription factors across diverse sets of epigenetic modifications, in any organism for which suitable data exist. The existence of specific transcription factor binding motifs whose recognition is driven by cytosine modifications may explain why transcription factors bind specific repetitive element loci, as opposed to every genome-wide iteration of the motif. Our work provides an initial foundation towards a better understanding of this important aspect of motif specificity.

## Availability

Cytomod is available at: https://github.com/hoffmangroup/cytomod. Persistent availability is ensured by [Zenodo](https://zenodo.org/), in which we have deposited the version of our code we used (https://doi.org/10.5281/zenodo.6345378). We also provide additional source code, containing other analysis scripts (https://github.com/hoffmangroup/2022modTFBSs; also archived at https://doi.org/10.5281/zenodo.6347792). All source code is licensed under a [GNU General Public License, version 3 (GPLv3)](https://www.gnu.org/licenses/gpl-3.0.en.html), except for CentriMo, which retains its original license. We have additionally archived our full set of scores for every assessed hypothesis pair (https://doi.org/10.5281/zenodo.6345400). We have deposited all CUT&RUN sequencing data and peak calls generated for this work in GEO (GSE198458).

## Authors’ contributions

Conceptualization, M.M.H.;Data Curation, C.V., N.J.W., and H.S.;Formal Analysis, C.V. and T.L.B.; Investigation, C.V. and C.A.I.; Methodology, C.V., T.L.B., and M.M.H.; Software, C.V., J.J., T.L.B.; Visualization, C.V; Validation, C.V., C.A.I., S.Y.S., S.M.L., and S.J.H.; Writing — Original Draft, C.V., C.A.I., S.M.L, and S.J.H.; Writing — Review & Editing, C.V., C.A.I., M.K.S.-H., S.Y.S., D.J.A., A.C.F.-S., D.D.De C., S.J.H., T.L.B., and M.M.H.;Resources, M.K.S.-H., D.J.A., A.C.F.-S., D.D.De C., M.M.H.;Funding Acquisition, M.M.H.;Project Administration, C.V. and M.M.H.;Supervision, M.M.H.

## Acknowledgements

We thank William Stafford Noble and Charles E. Grant for useful discussions and contributions to the MEME Suite. We thank Mehran Karimzadeh for providing us with the K562 subset of his processed Human ENCODE ChIP-seq peak calls. We thank Andrew D. Smith, Meng Zhou Benjamin E. Decato and Egor Dolzhenko for their work on MethPipe^83,85^ and for rapidly working to clarify and address any issues. We thank Jaime Castro-Mondragón and Jacques van Helden for their work on RSAT matrix-clustering^115^ and for their assistance with modifications to permit clustering in an expanded-alphabet context. We thank Michael L. Waskom for his visualization work on the Seaborn^104^ Python package and for actively providing support. We thank Nicholas Khuu for assistance with library preparation and DNA sequencing. We thank Neil Weingarden for coordination and consultation regarding DNA sequencing. We thank Carl Virtanen and Zhibin Lu for technical assistance. This research was enabled by support provided by: [Globus](https://www.globus.org/),^159,160^ [Compute Canada](https://www.computecanada.ca/) (specifically, [WestGrid](https://www.westgrid.ca/), [SHARCNET](https://www.sharcnet.ca/), and [SciNet](https://www.scinethpc.ca/)^161^), and the Bioinformatics and HPC Core, University Health Network. We thank [Life Science Editors](https://www.lifescienceeditors.com/) for editing services.

