## Supplementary material for "Modeling methyl-sensitive transcription factor motifs with an expanded epigenetic alphabet": Figures S1-S6; Appendix A; Table S1

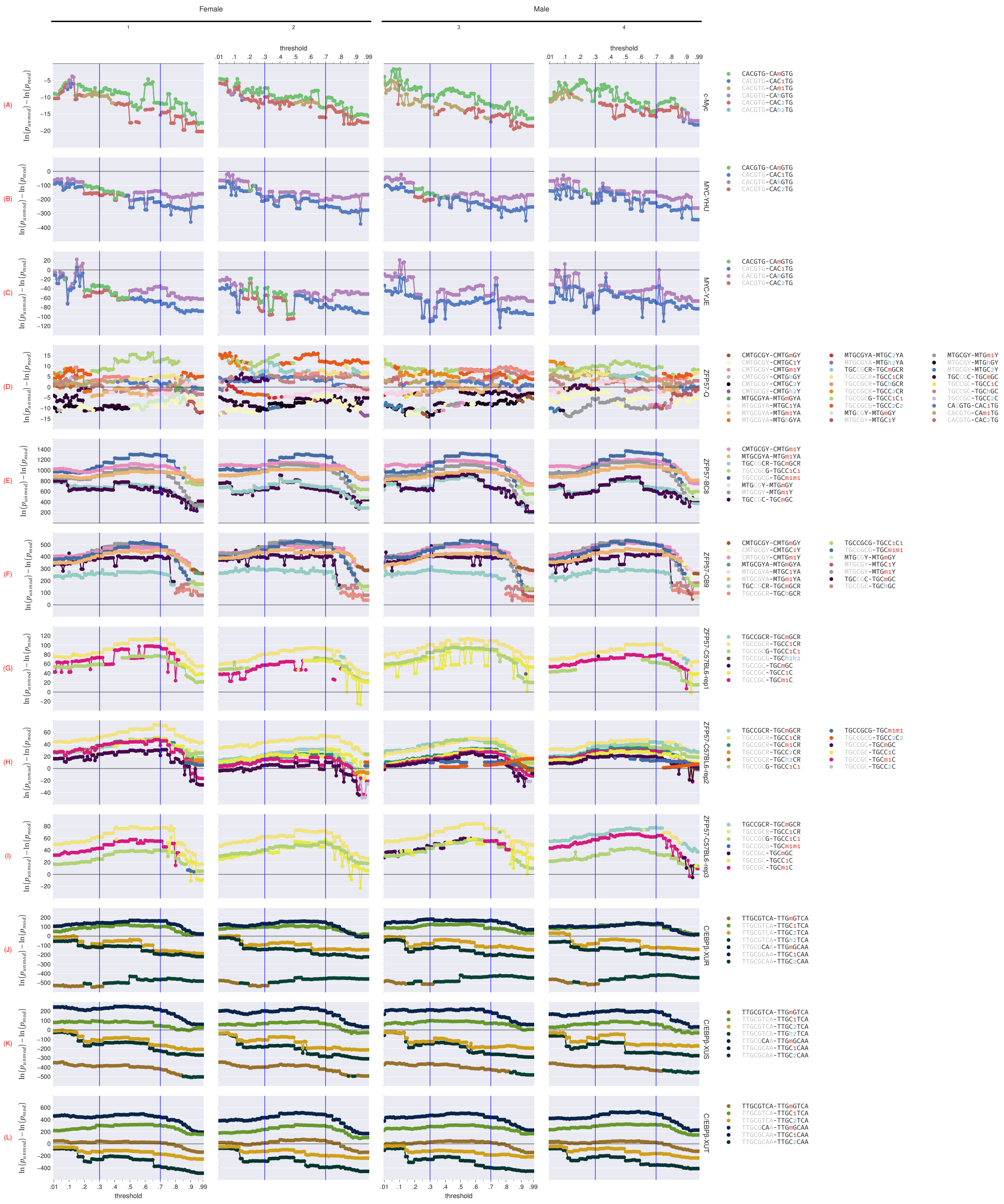

Female
Male
1
2
3
4
threshold
(A)
(B)
(C)
(D)
(E)
(F)
(G)
(H)
(I)
(J)
(K)
(L)
c-Myc
MYC-YHU
MYC-YJE
ZFP57-Q
ZFP57-BC8
ZFP57-CB9
ZFP57-C57BL6-rep1
ZFP57-C57BL6-rep2
ZFP57-C57BL6-rep3
C/EBPβ-XUR
C/EBPβ-XUS
C/EBPβ-XUT

**Figure S1 (preceding page). Relationship between unmodified versus modified motif statistical significance of central enrichment (from CentriMo[62]) and modified base calling thresholds across different whole genome bisulfite sequencing (WGBS) and oxidativeWGBS (oxWGBS) specimens, in mice.[59]** We compared each unmodified motif, at each threshold, to its top three most significant modifications for c-Myc, C/EBPβ, and replicate 2 of the C57BL/6 ZFP57 samples, but only the single most significant modification for all other ZFP57 samples. The displayed motif pairs changes at individual thresholds, depending on which motif pairs stay in the top three. We called all ZFP57 peaks, excepting Quenneville et al.[37], using a stringent MACS[68] threshold of $q = 0.00001$. Sign of value indicates preference for the unmodified (negative) motif or the modified (positive) motif. Rows: single chromatin immunoprecipitation-sequencing (ChIP-seq) replicates for a particular transcription factor target, consisting of: 3 c-Myc (Krepelova et al.[65], `ENCFF001YHU`, and `ENCFF001YJE`), 6 ZFP57 (from Quenneville et al.[37], and BC8 and CB9 cells, from Strogantsev et al.[38]), and 3 C/EBPβ (`ENCFF001XUR`, `ENCFF001XUS`, and `ENCFF001XUT`). Columns: replicates of WGBS and oxWGBS. We additionally depict C/EBPβ alone in Figure S3.

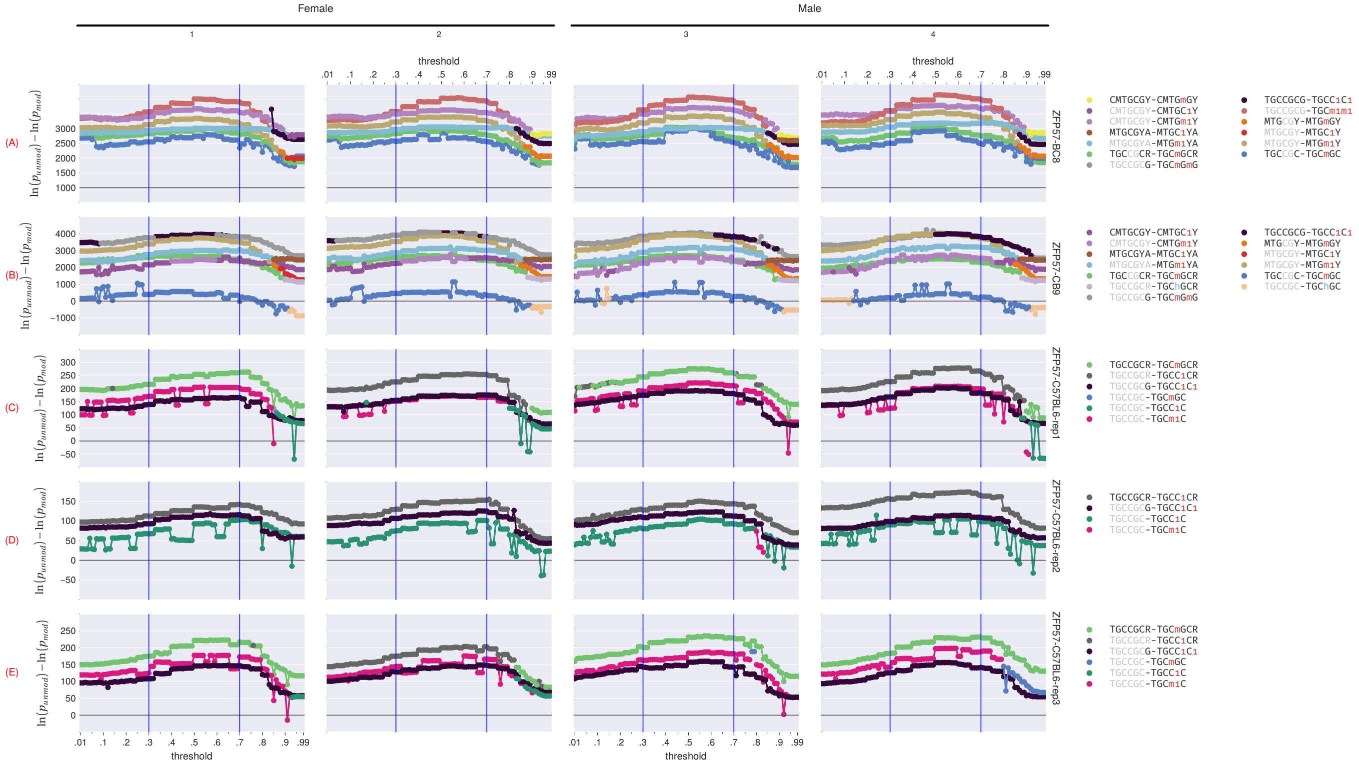


**Figure S2. Relationship between unmodified versus modified ZFP57 statistical significance of central enrichment (from CentriMo[62]) and modified base calling thresholds across different WGBS and oxWGBS specimens, in mice.[59]** We compared each unmodified motif, at each threshold, to its top most significant modification. The displayed motif pairs changes at individual thresholds, depending on which motif pairs stay at the top. We called all ZFP57 peaks using the default MACS[68] threshold of $q = 0.05$. Sign of value indicates preference for the unmodified (negative) motif or the modified (positive) motif. Rows: single ChIP-seq replicates for a particular transcription factor target, consisting of all ZFP57 replicates (from Quenneville et al.[37], and BC8 and CB9 cells, from Strogantsev et al.[38]). Columns: replicates of WGBS and oxWGBS.

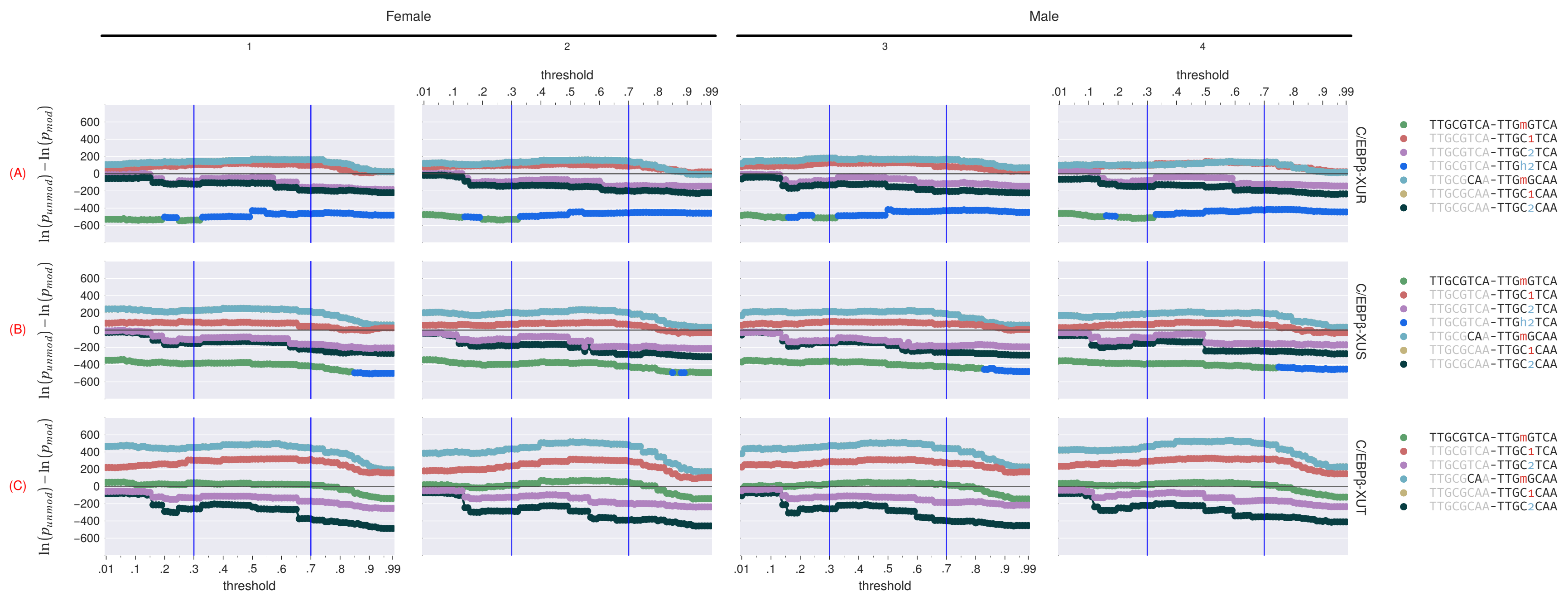


**Figure S3. Relationship between unmodified versus modified C/EBPβ statistical significance of central enrichment (from CentriMo[62]) and modified base calling thresholds across different WGBS and oxWGBS specimens, in mice.[59]** We compared each unmodified motif, at each threshold, to its top three most significant modifications. The displayed motif pairs changes at individual thresholds, depending on which motif pairs stay in the top three. Sign of value indicates preference for the unmodified (negative) motif or the modified (positive) motif. Rows: single ChIP-seq replicates for a particular transcription factor target, consisting of all C/EBPβ replicates (`ENCFF001XUR`, `ENCFF001XUS`, and `ENCFF001XUT`). Columns: replicates of WGBS and oxWGBS. We additionally depict all of our other tested transcription factors in Figure S1.

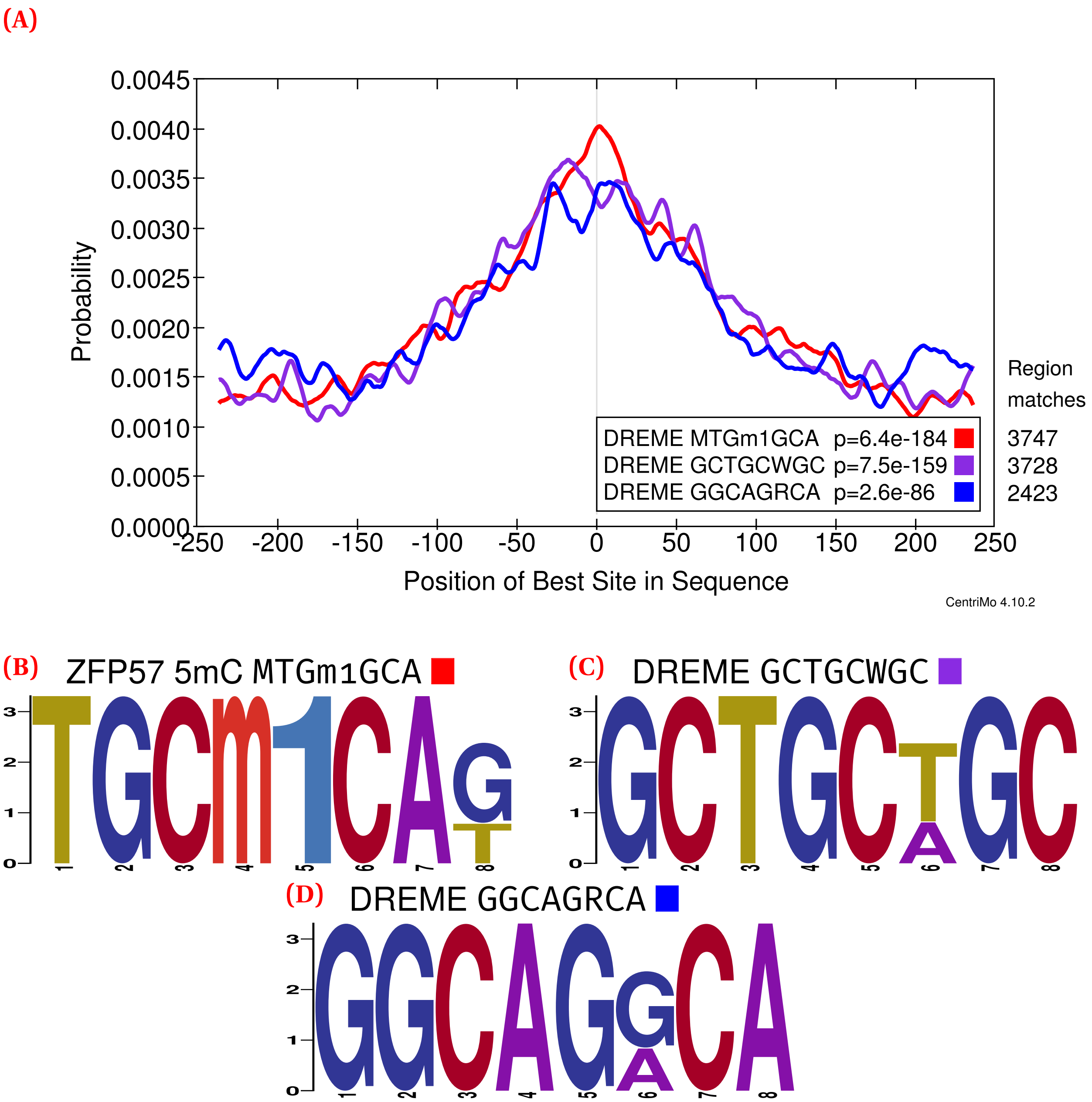


**Figure S4.** **ZFP57 (Strogantsev et al.[38] CB9; 56 142 ChIP-seq peaks) CentriMo analysis of *de novo* and JASPAR motifs (Methods).** Listed p-values computed by CentriMo.[62] Depicts female replicate 2 of the combined WGBS and oxWGBS data[59] at a 0.7 modification threshold. (A) the CentriMo result with an expected ZFP57 methylated motif (red), top DREME unmodified motif (purple), and second DREME unmodified motif (blue). (B) Sequence logo of the reverse complement of the CentriMo result with an expected ZFP57 methylated motif. (C) Sequence logo of the top DREME unmodified motif. (D) Sequence logo of the second DREME unmodified motif.

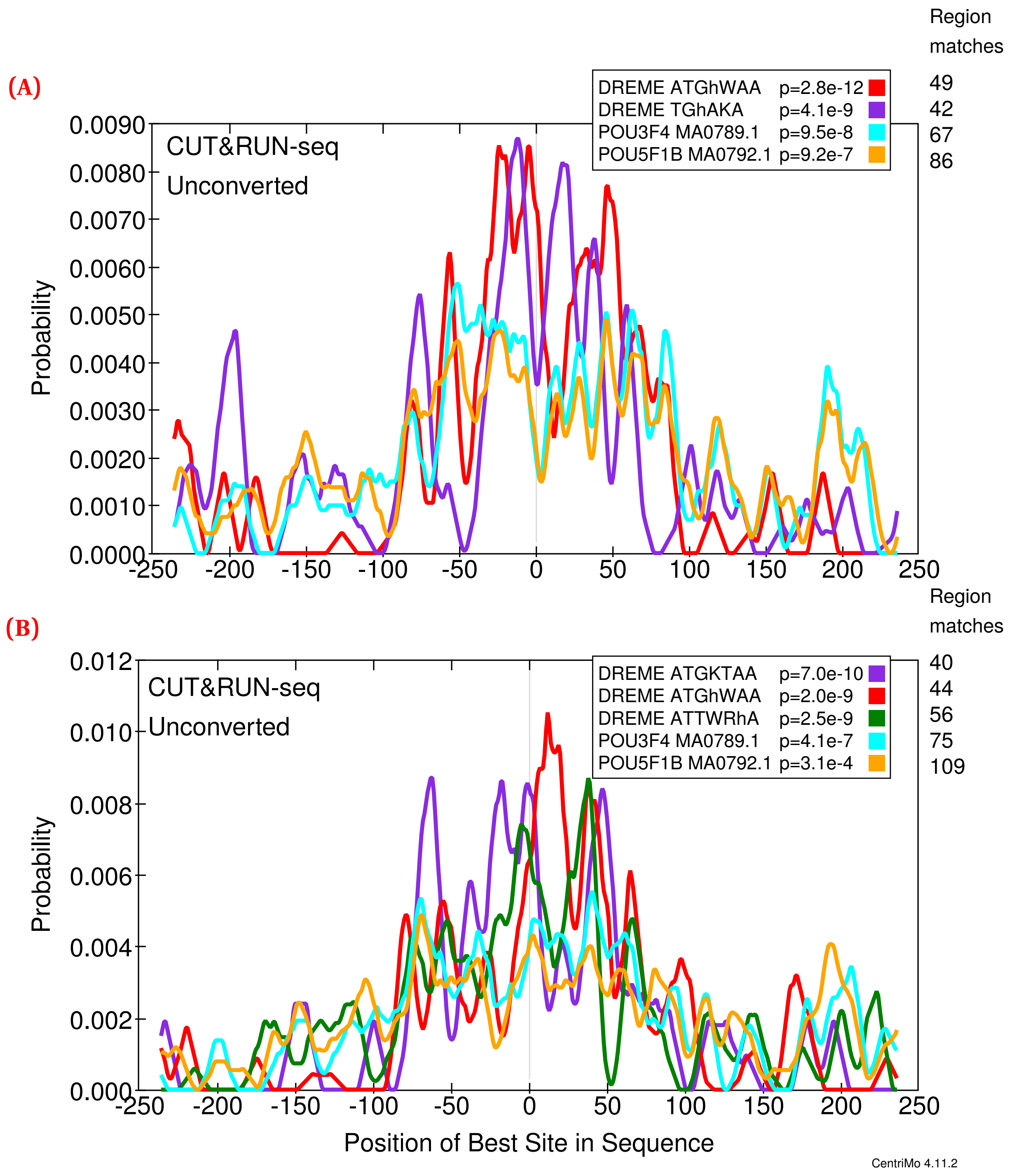
(A)
Region
matches
DREME ATGhWAA p=2.8e-12 49
DREME TGhAKA p=4.1e-9 42
POU3F4 MA0789.1 p=9.5e-8 67
POU5F1B MA0792.1 p=9.2e-7 86
CUT&RUN-seq
Unconverted
Probability
0.0090
0.0080
0.0070
0.0060
0.0050
0.0040
0.0030
0.0020
0.0010
0.0000
-250 -200 -150 -100 -50 0 50 100 150 200 250
(B)
Region
matches
DREME ATGKTAA p=7.0e-10 40
DREME ATGhWAA p=2.0e-9 44
DREME ATTWRhA p=2.5e-9 56
POU3F4 MA0789.1 p=4.1e-7 75
POU5F1B MA0792.1 p=3.1e-4 109
CUT&RUN-seq
Unconverted
Probability
0.012
0.010
0.008
0.006
0.004
0.002
0.000
-250 -200 -150 -100 -50 0 50 100 150 200 250
Position of Best Site in Sequence
CentriMo 4.11.2

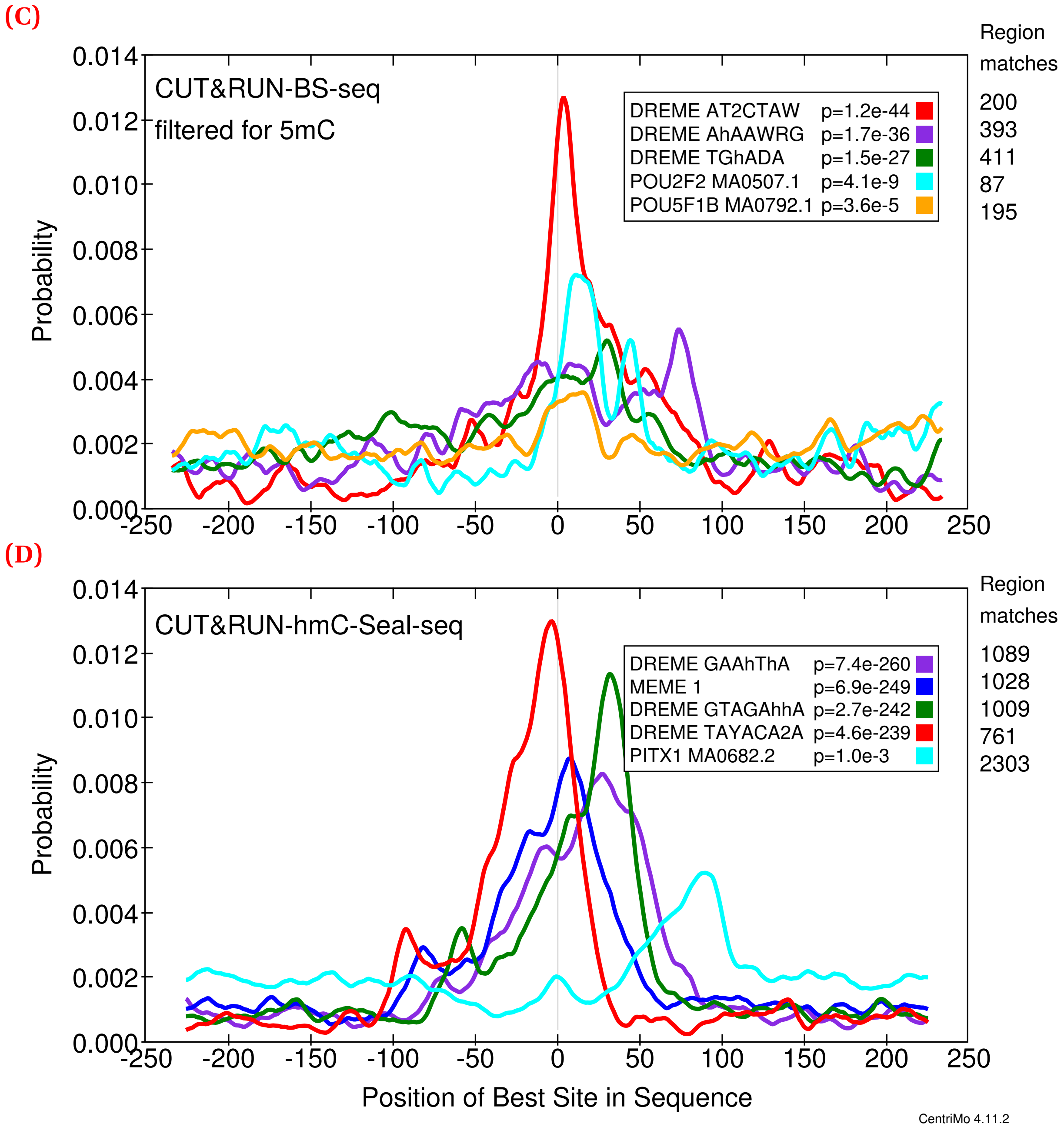


**Figure S5.** **CentriMo[62] results for OCT4 cleavage under targets and release using nuclease (CUT&RUN) in mouse embryonic stem cells (mESCs).** Motifs include the top three DREME motifs with colour indicating rank: first (red); second (purple), third, where applicable (dark green). Motifs also include the top non-POU5F1 JASPAR motif (cyan), and the JASPAR POU5F1B motif (orange). Both of these motifs come from the JASPAR 2020[67] core vertebrate set. We generated these results using 500 bp regions centred upon the summits of MACS 2[68] peaks generated from those CUT&RUN fragments ≤120 bp. We called peaks using IgG controls and without any spike-in calibration (Methods). Listed p-values computed by CentriMo.[62] For consistency, we depict the JASPAR sequence logo using MEME's relative entropy calculation and colouring. Also depicts the top MEME[69] *de novo* motif (blue). We depict the first replicates of 5-methylcytosine (5mC) and 5-hydroxymethylcytosine (5hmC) data in Figure 6. (A) replicate 1 unconverted (227 CUT&RUN peaks) sequences. (B) replicate 2 unconverted (265 CUT&RUN peaks) sequences. (C) replicate 2 bisulfite-converted (methylated; 2077 CUT&RUN peaks) sequences. (D) replicate 2 hmC-Seal (11 615 CUT&RUN peaks) sequences.

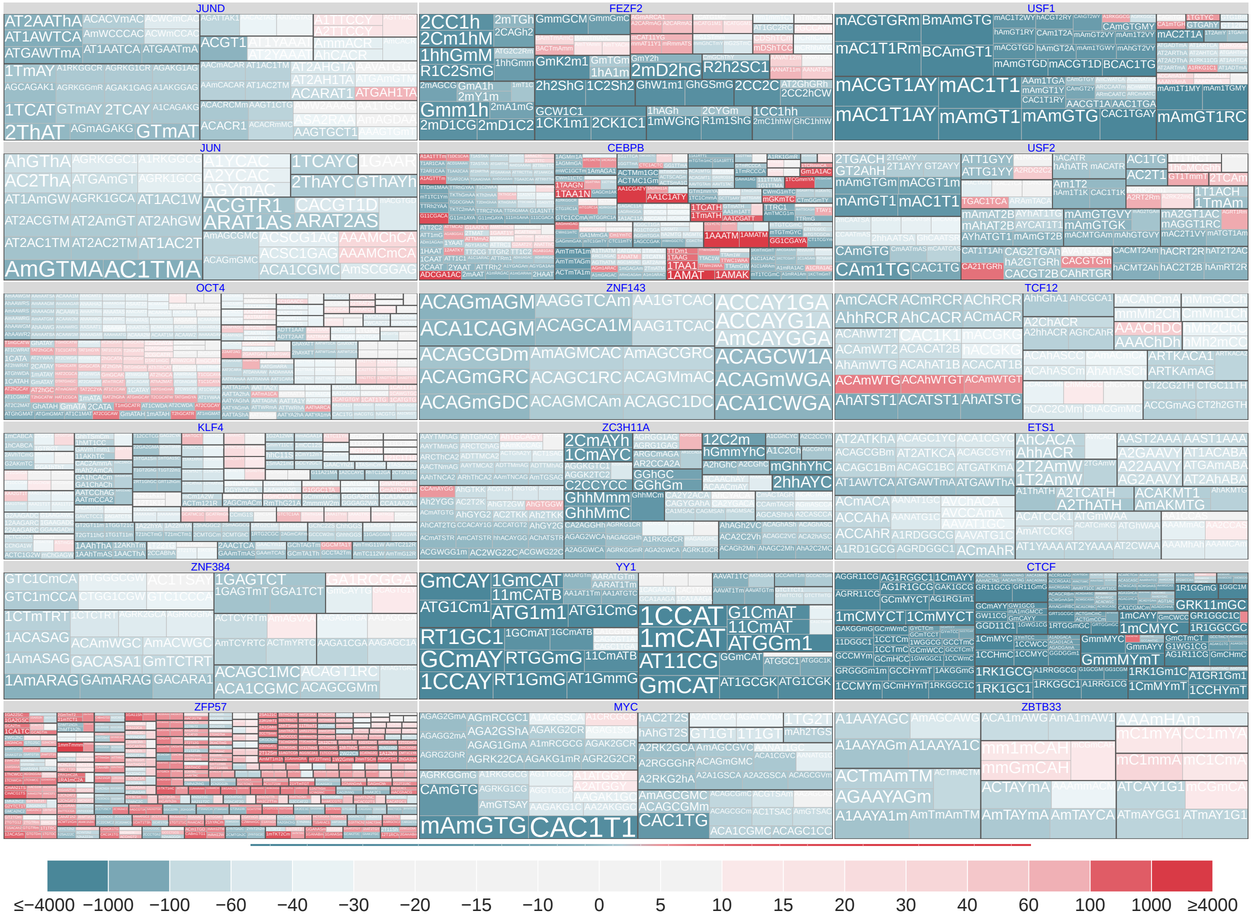


**Figure S6. Modified versus unmodified motifs, combining score and cluster information, for a wide array of transcription factors.** These plots come from non–spike-in calibrated data, for the 500 bp regions surrounding peak summits. We clipped scores above or below ±4000 and plotted them at a threshold that maximizes dynamic range where most scores occur. Some combinations of the displayed hypothesis pairs have multiple data points. For example, these points can correspond to multiple identical hypothesis pairs, but for different data sub-types or stringencies. We aggregated these multiple data points by plotting the maximum score. An asymmetric, diverging, colour scale further highlights modification-preferring motifs.

### Appendix A Recommendations for modified nucleobase nomenclature

Interest in different covalent DNA modifications and improvements in sequencing technologies have created a greater need for computational analyses of modified sequence data. To encourage standardization, we recommend symbols for various modified nucleobases (Table S1). We use lowercase letters (`a`–`z`) for specific nucleobase forms and numerals (`0`–`9`) to specify complements without any information loss. The list does not provide symbols for all DNA bases found in nature, but may provide guidance for those who need to select symbols for these bases.

The list here reserves specific symbols in an attempt to reduce contradictory definitions. We reserved all uppercase letters of the Latin alphabet for allocation by the International Union of Pure and Applied Chemistry (IUPAC), in addition to those already specified.[70] We use lowercase letters, and there do not exist sufficient unassigned letters to restrict ourselves to those without a different meaning in uppercase. Many applications, including the MEME Suite, only support Latin letters and numerals.

We recommend referring to any covalent modification at atomic position ⟨*position*⟩, specific modification ⟨*modification*⟩, and original nucleobase ⟨*base*⟩ as ⟨*position*⟩⟨*modification*⟩⟨*base*⟩. For example, this leads to "5mC" as the abbreviation for 5-methylcytosine. In particular, we recommend using no punctuation to demarcate the position from the modification and placing the numeral before the base modified. For example, others have occasionally abbreviated 6-methyladenine as "m6A", but we recommend the use of "6mA" instead when the modification occurs in DNA rather than in RNA.[165]

We incorporated the core symbols for cytosine modifications (Table 1) into our broader recommendations Table S1. While we specified a set of ambiguity codes for our usage here (Table 2), we do not recommend general definitions. Instead, we suggest reserving (1) the end of the lowercase Latin alphabet and (2) numerals. Avoiding universal assignment of these codes makes it more likely that the 62 symbols of the alphanumeric alphabet will prove sufficient for a variety of uses. As implemented here, we recommend assigning ambiguity codes starting from the end of the Latin alphabet or set of numerals, beginning with the most equivocal ambiguity code (such as `z` or `9`).

| Nucleobase | | | Complement | |
|---|---|---|---|---|
| Abbreviation | Name | Symbol | Name | Symbol |
| Covalent modifications of cytosine | | | | |
| 5mC | 5-methylcytosine | m | guanine:5mC | 1 |
| 5hmC | 5-hydroxymethylcytosine | h | guanine:5hmC | 2 |
| 5fC | 5-formylmethylcytosine | f | guanine:5fC | 3 |
| 5caC | 5-carboxylmethylcytosine | c | guanine:5caC | 4 |
| Covalent modifications of thymine | | | | |
| 5hmU | 5-hydroxymethyluracil | g | adenine:5hmU | |
| 5fU | 5-formyluracil | e | adenine:5fU | |
| 5caU | 5-carboxyluracil | b | adenine:5caU | |
| Covalent modifications of adenine | | | | |
| 6mA | 6-methyladenine | a | thymine:6mA | |
| Covalent modifications of guanine | | | | |
| 8-oxoG | 8-oxoguanine | o | adenine:8-oxoG (mismatch) | |
| Reserved synthetic bases | | | | |
| Xao | xanthosine | n | | |

**Table S1. Recommendations for the nomenclature of modified nucleobases, grouped by the unmodified nucleobase.** Numeral symbols for complements allow their differentiation from bases complementary to unmodified forms. One could reassign these numerals when analyzing other sets of covalent modifications. Since xanthosine can base-pair with multiple bases,[166] we list no complement for it.
